# A new gene finding tool GeneMark-ETP significantly improves the accuracy of automatic annotation of large eukaryotic genomes

**DOI:** 10.1101/2023.01.13.524024

**Authors:** Tomas Bruna, Alexandre Lomsadze, Mark Borodovsky

## Abstract

Large-scale genomic initiatives, such as the Earth BioGenome Project, require efficient methods for eukaryotic genome annotation. Here we present an automatic gene finder, GeneMark-ETP, integrating genomic-, transcriptomic- and protein-derived evidence that has been developed with a focus on large plant and animal genomes. GeneMark-ETP first identifies genomic loci where extrinsic data is sufficient for making gene predictions with ‘high confidence’. The genes situated in the genomic space between the high confidence genes are predicted in the next stage. The set of high confidence genes serves as an initial training set for the statistical model. Further on, the model parameters are iteratively updated in the rounds of gene prediction and parameter re-estimation. Upon reaching convergence, GeneMark-ETP makes the final predictions and delivers the whole complement of predicted genes. GeneMark-ETP outperformed gene finders using a single type of extrinsic evidence. Comparisons with gene finders utilizing both transcript- and protein-derived extrinsic evidence, MAKER2, and TSEBRA, demonstrated that GeneMark-ETP delivered state-of-the-art gene prediction accuracy with the margin of outperforming existing approaches increasing in its applications to larger and more complex eukaryotic genomes.

## Introduction

Massive sequencing of eukaryotic genomes, e.g., the Earth BioGenome Project (Lewin et al. 2022), should be complemented with fast and accurate automatic tools of genome annotation. The development of such tools remains an active area of research. One of the long-standing challenges is to achieve optimal integration of the intrinsic evidence of protein-encoding by a nucleotide sequence with the extrinsic evidence of gene presence provided by endogenous RNA or known cross-species protein sequences. Gene finding methods developed before transition to massive genomic sequencing powered by NGS have been focused on using intrinsic evidence in the form of the Markov chain models reflecting the *k*-mer frequency patterns, probabilistic models of functional sites controlling translation and splicing, intron/exon length distributions, etc. All these probabilistic models served as elements of a generalized HMM (GHMM), e.g., Genie (Kulp et al. 1996), GENSCAN (Burge and Karlin 1997), GeneID (Parra et al. 2000). The bottleneck of the application of the early methods was a tedious preparation of the supervised species-specific training sets. The training automation was first made in GeneMark-ES, an *ab initio* gene finder implementing a hierarchical procedure of iterative unsupervised training (Lomsadze et al. 2005; Ter-Hovhannisyan et al. 2008). The accuracy of gene predictions made by GeneMark-ES in compact fungal and protist genomes has been high and is still difficult to improve even by adding currently available extrinsic evidence. Since the advent of NGS, the rapid accumulation of RNA and protein information has opened ample opportunities for addition of extrinsic evidence (Guigo et al. 2006; Coghlan et al. 2008; Goodswen et al. 2012; Scalzitti et al. 2020). This addition proved to be critically important for increasing gene prediction accuracy in large plant and animal genomes having low gene density. The accuracy improvement of fully automated methods is a two-fold problem, it concerns improvements of both training and prediction steps.

One kind of extrinsic evidence comes from known cross-species proteins. These proteins may guide predictions of the genes encoding proteins from the same family in a novel genome. Confident spliced alignment of a cross-species protein to a locus in the novel genome gives excellent starting point for delineation of the full exon-intron structure of the true gene. Spliced alignments have been used in several gene finders, e.g., in exonerate (Slater and Birney 2005), GenomeThreader (Gremme et al. 2005), and ProSplign (Kiryutin et al. 2007). Sequencing of expressed endogenous RNA provides another type of external evidence. Cufflinks (Trapnell et al. 2010), StringTie (Pertea et al. 2015; Kovaka et al. 2019), PsiCLASS (Song et al. 2019), etc. map short RNA-seq reads to genome to identify exon/intron borders of the genes having detectable levels of expression. Notably, a method leveraging any type of extrinsic evidence is useful only for finding a subset of genes for which such evidence exists.

The goal of improving integration of intrinsic and extrinsic evidence has continued to motivate creation of new gene finding methods for more than two decades, e.g., GAZE (Howe et al. 2002), Combiner (Allen et al. 2004), JIGSAW (Allen and Salzberg 2005), Evigan (Liu et al. 2008), EVidenceModeler (Haas et al. 2008), MAKER2 (Holt and Yandell 2011), IPred (Zickmann and Renard 2015), GeMoMa (Keilwagen et al. 2018), LoReAn (Cook et al. 2019), GAAP (Kong et al. 2019), and FINDER (Banerjee et al. 2021), BRAKER1 (Hoff et al. 2016), BRAKER2 (Bruna et al. 2021) and TSEBRA (Gabriel et al. 2021).

GeneMark-ETP takes several steps of integration. First, GeneMarkS-T, the GHMM based *ab initio* gene finder for transcripts (Tang et al. 2015) is used to predict protein-coding regions (CDSs) in transcripts assembled from RNA-seq reads by StringTie2 (Kovaka et al. 2019). Second, the predicted CDSs are validated or modified, if the protein level evidence suggests so, by the module that we call GeneMarkS-TP. Next, the CDSs predicted in transcripts are mapped to the genome. We show that the resulting gene models have distinctly high Precision and call them *high-confidence genes (HC genes)*. The set of HC genes is used as an initial training set for the genomic GHMM used by GeneMark-ETP to predict genes situated in genomic regions between the HC genes. The extrinsic information (hints from spliced aligned proteins and mapped RNA-seq reads) is plugged in at this step into the Viterbi algorithm. The rounds of genome wide gene prediction and model parameters estimation continue until convergence.

For benchmarking, we selected seven eukaryotic genomes, both GC-homogeneous and GC-inhomogeneous: *Arabidopsis thaliana*, *Caenorhabditis elegans, Drosophila melanogaster, Solanum lycopersicum*, *Danio rerio*, *Gallus gallus*, and *Mus musculus*. Along with GeneMark-ETP we tested GeneMark-ET, GeneMark-EP+, the pipelines BRAKER1, BRAKER2, MAKER2, as well as TSEBRA. The tests demonstrated the state-of-the-art performance of GeneMark-ETP, whose margin of improvement over other gene finders was increasing with the increase in the genome length.

## Results

### Introductory comments

Here, we introduce definitions of several terms used throughout the text. The goal of gene finding algorithms is to predict protein-coding genes which may have several alternative transcripts. Upon RNA transcripts assembly (by StringTie2), transcripts representing one and the same gene (alternative RNA transcripts) are determined. The alternative mRNA transcripts may have identical CDS sequences but may differ in the UTR regions. We are concerned only with the protein (CDS) isoforms.

We measure gene prediction accuracy at two levels, the *gene level* and the *exon level*. A protein-coding gene prediction is considered to be correct if *at least one* of the predicted CDS isoforms matches one of the annotated CDS isoforms. A protein-coding exon is predicted correctly if it matches exactly an annotated exon.

Most previous publications on gene finding methods and their benchmarking have used the term ‘Specificity’ for the measure defined as #*TP*/(#*TP* + #*FP*), where #*TP* is the number of true positives and #*FP* is the number of false positives. However, in the related and fast-growing field of machine learning, the term ‘Specificity’ has the meaning of true negative rate defined as #*TN*/(#*TN* + #*TP*), where #*TN* is the number of true negatives. To unify the designations, due to the request of the Genome Research Editors, we have switched here to the term ‘Precision’ (Pr) for #*TP*/(#*TP* + #*FP*).

Thus, we characterize the prediction accuracy (at a gene or an exon level) by Sensitivity, *Sn* = #*TP*/(#*TP* + #*FN*), and Precision, *Pr* = #*TP*/(#*TP* + #*FP*), where #TP and #FP are defined above, and #FN is the number of false negative predictions.

As a single accuracy measure, we use the F1 score, the harmonic mean of Sensitivity and Precision, *F*1 = 2 × *Sn* × *Pr*/(*Sn* + *Pr*) or *F*1 = #*TP*/(#*TP* + (#*FN* + #*FP*)/2). For convenience, the F1 score is multiplied by 100. Note that the latter formula for F1 is not equivalent to the former one if Sensitivity and Precision are computed using different sets of positive examples (annotated CDSs). Therefore, to avoid ambiguities, we use the former formula in all the computations.

### Integration of extrinsic evidence in GeneMarkS-TP

At the first step of the GeneMark-ETP pipeline (Fig. 1, Supplemental Figs. S1-S6), short RNA-seq reads were assembled into transcripts by StringTie2 (Kovaka et al. 2019). Next, iterative unsupervised training of GHHM of GeneMarkS-T was done on these assembled transcripts using the rounds of model parameters estimation and *ab initio* gene predictions (Tang et al. 2015).

**Figure 1.**
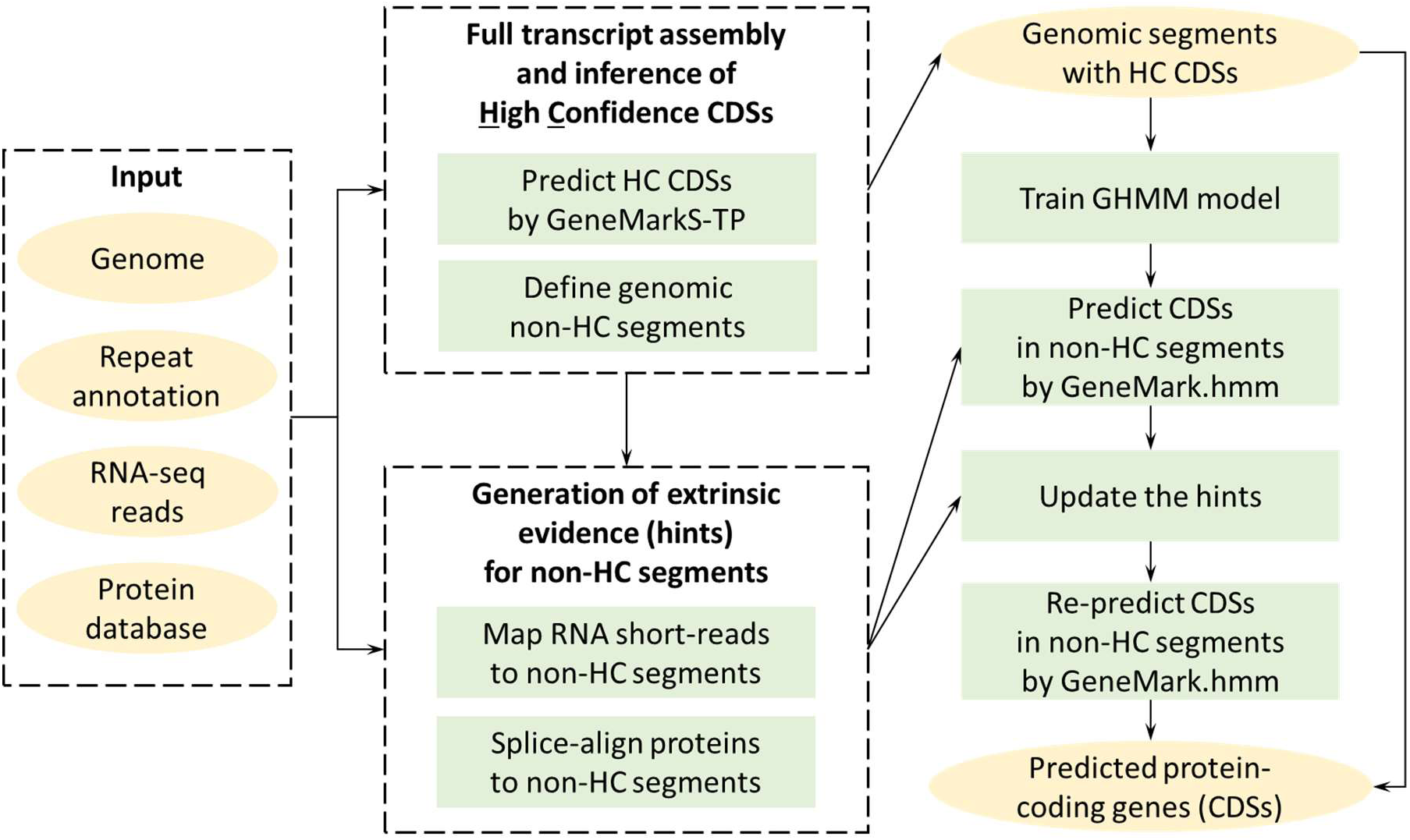
High level diagram of the GeneMark-ETP processing of genomic, RNA-seq, and protein sequences. For more details, see Fig. 2 and Supplemental Figs. S1-S6.

Running GeneMarkS-T is the first step in the GeneMarkS-TP module (Fig. 2). The CDSs predicted by GeneMarkS-T in assembled transcripts were translated, and the amino acid sequences were aligned with homologous cross-species proteins found by a database search. The alignments served as guides for corrections of the initially predicted CDSs (see Methods). The magnitudes of changes in Sn and Pr upon the corrections depended on the size of the protein database used by GeneMarkS-TP (Table 1). Corrections in initially predicted CDSs increased the Precision values, on average, by 25 percentage points, reaching close to or even higher than 90%. Thus, the refined CDSs were dubbed High Confidence CDSs (HC CDSs) (Table 1 and Supplemental Table S1).

**Figure 2.**
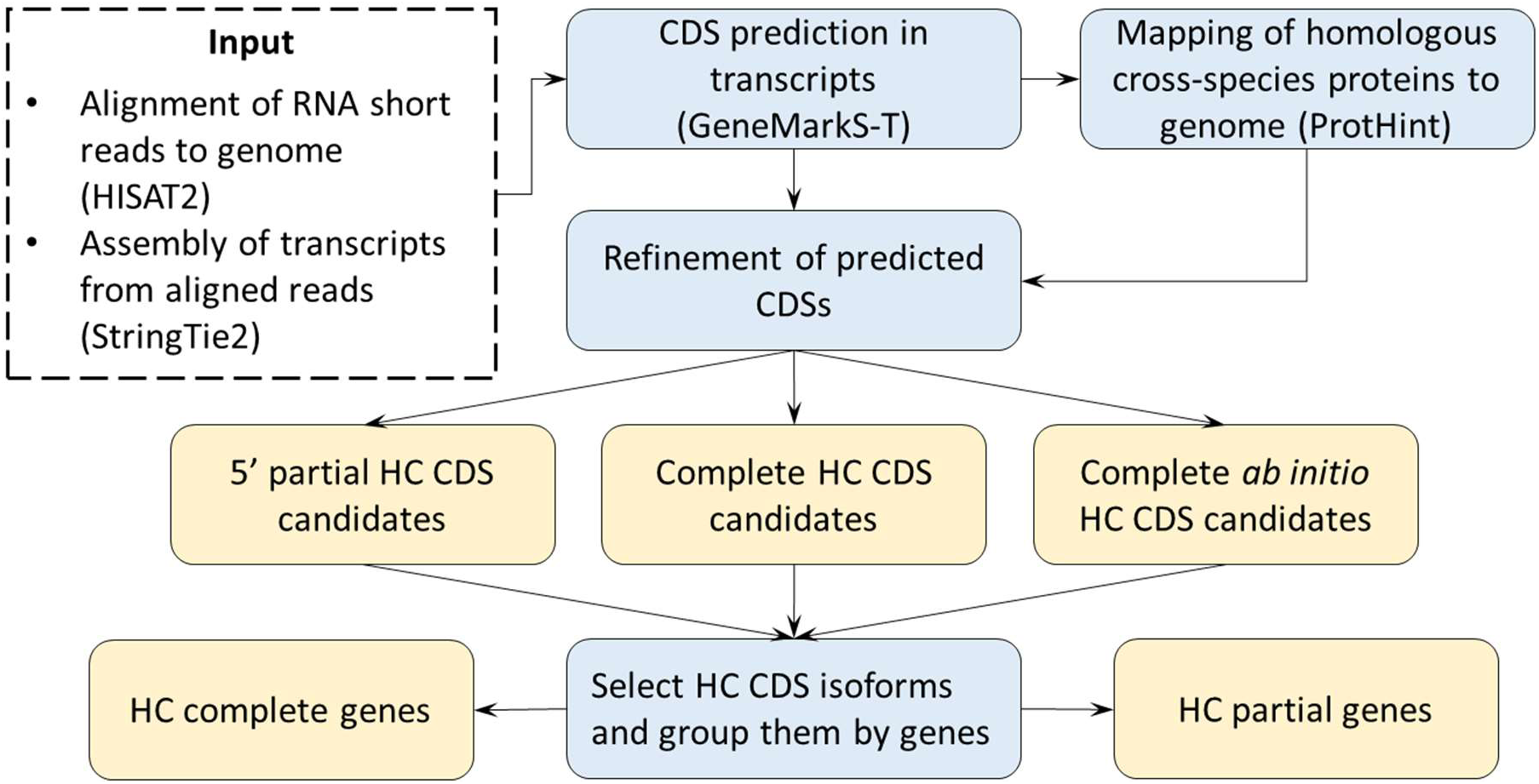
The diagram illustrates the work of the GeneMarkS-TP module generating the CDSs and genes predicted with high confidence (see also Supplemental Figs. S1-S3.)

**Table 1.** Statistics characterizing the GeneMarkS-T *initial gene predictions* made in assembled transcripts as well as the HC genes predicted by the GeneMarkS-TP module. Two versions of a reference protein database were used for each species: the ‘Species excluded’, as well as the ‘Order excluded’ (see Data Sets). Refinement made by GeneMarkS-TP reduced the number of initial predictions and produced a significant increase in Pr. In most of the genomes there was also a positive change in Sn, especially in large inhomogeneous genomes of *G. gallus* and *M. musculus*. Additional data is provided in Supplemental Table S1.

| Species | # of genes initially predicted by GeneMarkS-T | Sn/Pr of GeneMarkS-T predicted genes | Order excluded DB |  | Species excluded DB |  |
| --- | --- | --- | --- | --- | --- | --- |
|  |  |  | # of HC genes | Sn/Pr of HC genes | # of HC genes | Sn/Pr of HC genes |
| *C. elegans* | 14,746 | 46.8 / 63.4 | 8,062 | 35.7 / 88.4 | 11,399 | 51.7 / 90.6 |
| *A. thaliana* | 17,589 | 51.2 / 79.9 | 16,008 | 56.7 / 97.3 | 16,551 | 58.8 / 97.6 |
| *D. melanogaster* | 10,163 | 59.6 / 81.8 | 8,109 | 55.0 / 94.7 | 9,223 | 63.7 / 96.3 |
| *S. lycopersicum* | 19,526 | 68.0 / 78.4 | 17,231 | 75.1 / 95.3 | 17,489 | 75.8 / 95.2 |
| *D. rerio* | 22,992 | 59.6 / 59.9 | 16,918 | 67.0 / 88.5 | 16,573 | 66.9 / 90.4 |
| *G. gallus* | 17,381 | 49.6 / 47.0 | 12,473 | 74.4 / 89.1 | 12,564 | 74.0 / 88.4 |
| *M. musculus* | 15,819 | 49.6 / 63.2 | 13,057 | 63.5 / 93.2 | 12,965 | 63.9 / 94.5 |

We saw that both the numbers of GeneMarkS-T predictions (the HC CDS candidates) and the numbers of HC CDSs predicted by GeneMarkS-TP correlated with the numbers of the RefSeq annotated CDSs (Table 1, Supplemental Fig. S7). The HC CDSs could give accurate predictions of exon-intron structure coordinates for up to 60% of protein-coding genes present in the RefSeq genome annotations (Table 1).

The numbers of HC CDSs predicted with the larger ‘Species excluded’ databases did not differ significantly from the numbers of HC CDSs predicted with the use of the smaller ‘Order excluded’ databases (Table 1, Supplemental Fig. S7). Also, the Precision values of the HC CDSs did not change significantly upon transition from the ‘Order excluded’ to the ‘Species excluded’ database.

At the step preceding the generation of the final set of gene predictions, we have had a large set of gene candidates. The gene candidates could be divided into groups with different level of extrinsic support: (i) those with full extrinsic support having all the exon borders supported by high scoring hints; (ii) those with partial extrinsic support, having high scoring hints for some but not all exon borders; (iii) those that were predicted *ab initio* and had an *a posteriori* detected match to a low scoring hint of at least one exon border; (iv) those predicted *ab initio* and had no match to any extrinsic hints. Importantly, the HC CDSs belong to the first two categories. The meaning of an ‘*a posterior*i detected match to low scoring hints’ is that such hints were not used in the gene prediction algorithm.

Now, to focus solely on results, we must skip further details related to the description of the pipeline (see Methods, Figs. 1-2, and Supplemental Figs. S1-S6).

One could see that for all seven genomes, the Precision values of the candidate genes decreased significantly upon a decrease in the level of extrinsic support, thus indicating an increase in false positive rates (Table 2).

**Table 2.** Distribution of the gene prediction candidates among the four groups that differ by the level of extrinsic support. In the ‘Match to low scoring hints’ category we placed the candidates that were predicted fully *ab initio* while in a *posteriori* analysis a match of at least one exon border to a low scoring extrinsic hint was detected. The category ‘No extrinsic match’ contains the *ab initio* predictions that have no match to any extrinsic hints (see text). Average gene level Pr values are listed for each group. Protein databases are described in the section Data Sets.

| Species | Support for predicted gene candidates | Order excluded DB |  | Species excluded DB |  |
| --- | --- | --- | --- | --- | --- |
|  |  | # of candidates | Precision % | # of candidates | Precision % |
| *C. elegans* | Fully extrinsic | 7,676 | 88.9 | 10,778 | 91.6 |
|  | Partially extrinsic | 4,804 | 56.4 | 5,417 | 54.4 |
|  | Match to low scoring hints | 4,020 | 54.7 | 1,548 | 45.2 |
|  | No extrinsic match | 1,298 | 24.9 | 778 | 18.0 |
| *A. thaliana* | Fully extrinsic | 16,445 | 97.2 | 18,083 | 97.5 |
|  | Partially extrinsic | 4,825 | 64.4 | 5,807 | 55.7 |
|  | Match to low scoring hints | 1,794 | 50.2 | 1,360 | 30.1 |
|  | No extrinsic match | 2,964 | 27.9 | 1,128 | 9.4 |
| *D. melanogaster* | Fully extrinsic | 8,059 | 95.1 | 9,952 | 96.8 |
|  | Partially extrinsic | 2,328 | 49.3 | 2,751 | 44.9 |
|  | Match to low scoring hints | 1,043 | 57.1 | 165 | 44.9 |
|  | No extrinsic match | 1,369 | 41.6 | 377 | 15.9 |
| *S. lycopersicum* | Fully extrinsic | 17,639 | 95.2 | 18,420 | 95.0 |
|  | Partially extrinsic | 5,174 | 47.3 | 5,813 | 44.3 |
|  | Match to low scoring hints | 1,577 | 38.4 | 1,484 | 29.7 |
|  | No extrinsic match | 4,714 | 14.8 | 3,703 | 9.2 |
| *D. rerio* | Fully extrinsic | 15,691 | 89.8 | 15,501 | 92.6 |
|  | Partially extrinsic | 10,905 | 16.6 | 11,769 | 16.6 |
|  | Match to low scoring hints | 1,973 | 11.4 | 1,663 | 7.3 |
|  | No extrinsic match | 12,534 | 0.8 | 11,879 | 0.3 |
| *G. gallus* | Fully extrinsic | 11,856 | 89.3 | 11,547 | 89.9 |
|  | Partially extrinsic | 4,857 | 19.6 | 5,337 | 20.1 |
|  | Match to low scoring hints | 527 | 8.9 | 579 | 7.1 |
|  | No extrinsic match | 11,332 | 0.4 | 11,352 | 0.3 |
| *M. musculus* | Fully extrinsic | 13,556 | 94.6 | 13,769 | 96.2 |
|  | Partially extrinsic | 7,376 | 20.6 | 7,606 | 19.6 |
|  | Match to low scoring hints | 957 | 10.1 | 1,155 | 7.3 |
|  | No extrinsic match | 20,711 | 1.2 | 19,666 | 0.5 |

In the genomes of *D. rerio*, *G. gallus,* and *M. musculus*, having the largest average length of introns and intergenic regions among the seven genomes, the gene candidates of category *iv* had Pr values at the gene level below 1.5%, thus indicating an exceedingly high false positive rate. For these three large genomes as well as for the genome of *S. lycopersicum*, an exclusion of gene candidates of category *iv* led to 21 percentage points increase, on average, in the gene-level Pr, with a simultaneous 0.3 percentage points decrease in Sn (Supplemental Table S2). On the other hand, such a pruning of the gene candidates generated in the analysis of the three compact genomes, *A. thaliana*, *C. elegans*, and *D. melanogaster* would increase, on average, the gene level Pr by 3.7 percentage points and decrease Sn by 1.7 percentage points. These observations suggest that when sufficiently long eukaryotic genomes (e.g., genomes longer than 300 Mb) are considered, the gene candidates of category *iv* should not be included into the lists of predicted genes (CDSs).

Yet another observation is that the numbers of predicted genes are close to the numbers of annotated genes, while the total numbers of predicted CDSs (including alternative CDSs) are consistently and significantly smaller than those in annotation (Table 3). Indeed, alternative CDSs are predicted in relatively small numbers on the whole genome scale, since they are predicted only as CDS isoforms of the HC genes.

**Table 3.** Numbers of the predicted and annotated genes as well as individual CDSs (including all alternative CDSs). Note that the genomic CDSs and the corresponding transcript CDSs are supposed to be identical in sequence. The Order excluded reference databases were used by GeneMark-ETP (See Data Sets).

| Species | GeneMark-ETP |  | RefSeq annotation |  |
| --- | --- | --- | --- | --- |
|  | # of protein-coding genes | # of predicted genomic CDSs | # of protein-coding genes | # of annotated genomic CDSs |
| *C. elegans* | 18,820 | 19,806 | 19,969 | 28,544 |
| *A. thaliana* | 26,449 | 27,708 | 27,445 | 40,827 |
| *D. melanogaster* | 12,850 | 14,138 | 13,951 | 22,395 |
| *S. lycopersicum* | 24,420 | 26,341 | 25,158 | 31,911 |
| *D. rerio* | 28,608 | 31,961 | 25,610 | 42,929 |
| *G. gallus* | 17,275 | 21,433 | 17,279 | 38,534 |
| *M. musculus* | 23,956 | 27,686 | 22,405 | 58,318 |

### Assessment of the gene prediction accuracy

The accuracy of previously developed gene finding tools shows consistent decline with the increase in the length of eukaryotic genome. This pattern is due to the increasing difficulty of accurate finding of protein coding sequences constituting a diminishing fraction of the whole genome, while non-coding regions are taking larger and larger genomic space. Indeed, for compact genomes, the values of gene prediction Sn and Pr of the tools developed earlier and those developed later, including GeneMark-ETP, are relatively close and change incrementally (Fig. 3, Supplemental Fig. S8). However, for the large genomes, GeneMark-ETP makes significant improvements both for GC-homogenous *S. lycopersicum*, *D. rerio*, and, especially, for GC-inhomogeneous *G. gallus* and *M. musculus,* where we see the double-digit improvements of Sn and Pr (Fig. 4, Supplemental Fig. S8, Supplemental Tables S3-S4).

**Figure 3.**
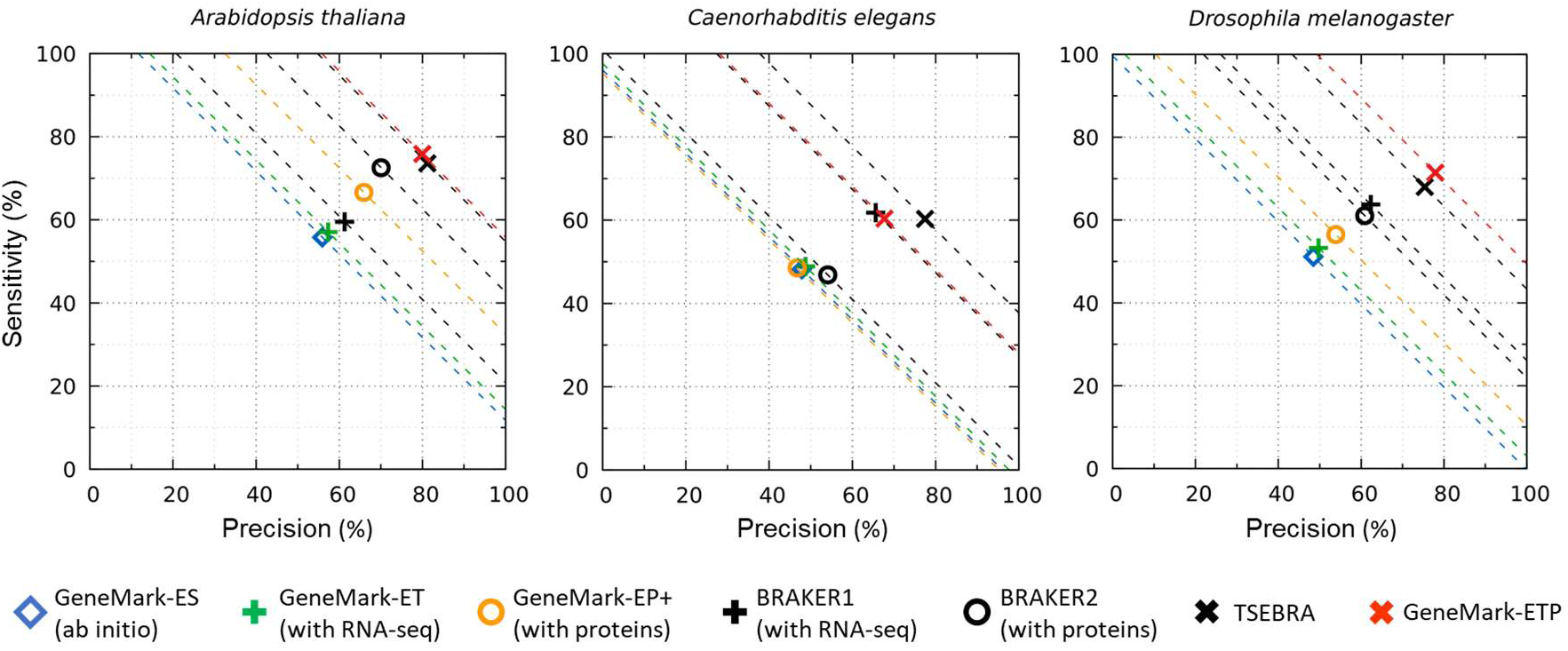
Gene level Sn and Pr of GeneMark-ETP and six other gene finders were computed for the three compact genomes: *C. elegans*, *A. thaliana*, and *D. melanogaster*. The dashed lines correspond to constant levels of (Sn+Pr)/2. The true positives were defined with respect to the complete set of annotated genes (either in RefSeq or Ensemble that had identical annotations of these genomes). The ‘Order excluded’ protein databases were used.

**Figure 4.**
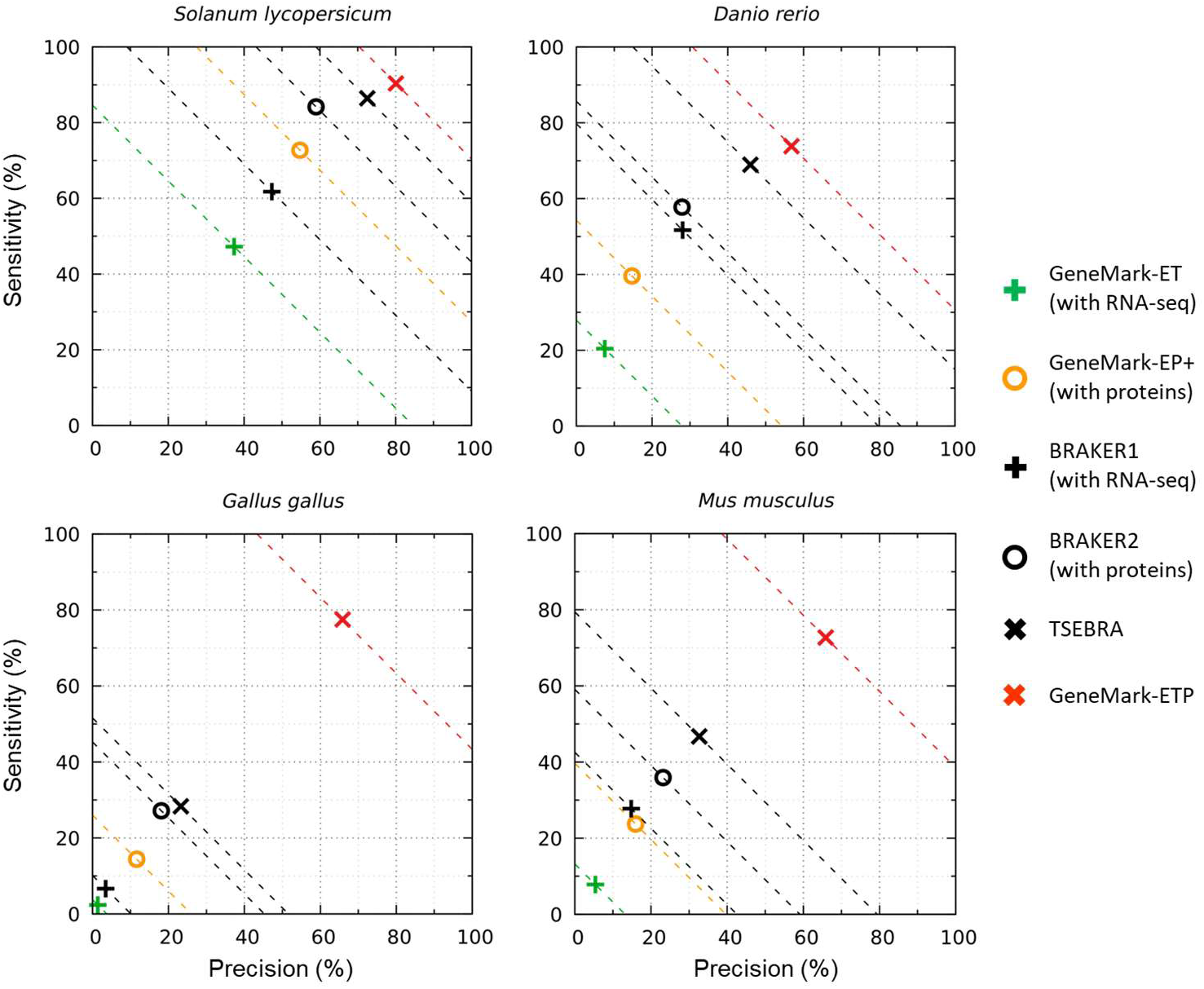
Gene level Sn and Pr of gene predictions made by GeneMark-ETP and five other gene finders in the four large genomes: *S. lycopersicum*, *D. rerio*, *G. gallus*, and *M. musculus*. The Sn and Pr values for each genome were defined with respect to subsets of genes present in the RefSeq and Ensemble genome annotations (the intersections and the unions of annotations, see text). For these genomes, gene candidates lacking extrinsic support did not make it into the GeneMark-ETP list of predicted genes (see text). The ‘Order excluded’ protein databases were used.

The results of comparison with the gene finders using a single type of extrinsic evidence were as follows. Over GeneMark-ET (utilizing RNA-seq reads mapping to augment the training process) the gene level F1 was improved, on average, by 19.6, 47.8, and 66.3 points for the groups of compact, large homogeneous, and large inhomogeneous genomes, respectively. Over GeneMark-EP+ (using spliced aligned cross-species proteins) the increases in F1 were, respectively, 14.2, 33.9, and 55.7 points, noticeably lower numbers than the ones related to GeneMark-ET (Supplemental Table S3). GeneMark-ETP also had shown improvements in F1 scores over BRAKER1 and BRAKER2, the pipelines containing GeneMark-ET and GeneMark-EP+, respectively. In comparison with BRAKER1, for the same three groups of genomes the increases in F1 were, on average, by 9.7, 29.2, and 58.5 points. In comparison with BRAKER2, the F1 average improvements were 11.2, 21.9, and 47.6 points. The better performance of BRAKER2 in comparison with BRAKER1 is in line with observed better performance of GeneMark-EP+ in comparison with GeneMark-ET, which are important parts of BRAKER2 and BRAKER1 respectively.

Next, we have considered gene predictions where two sources of extrinsic evidence are accounted for. First, three combinations of the sets of gene predictions made separately by GeneMark-ET and GeneMark-EP+ could be constructed (see Methods). Still, GeneMark-ETP outperformed even the ‘best’ combination of GeneMark-ET and GeneMark-EP+ gene predictions. For the *D. melanogaster* genome, taken as an example, GeneMark-ETP improved Sensitivity by 7.5 percentage points and Precision by 13 percentage points over the ‘best’ combination which Sn and Pr correspond to a ‘diamond’ point in Fig. 5. The gene level F1 scores were improved by GeneMark-ETP over the F1 scores of the ‘best’ combination of gene predictions by 10.1, 18.0, and 56.6 points for genomes of *D. melanogaster*, *S. lycopersicum*, and *G. gallus*, respectively (data not shown).

**Figure 5.**
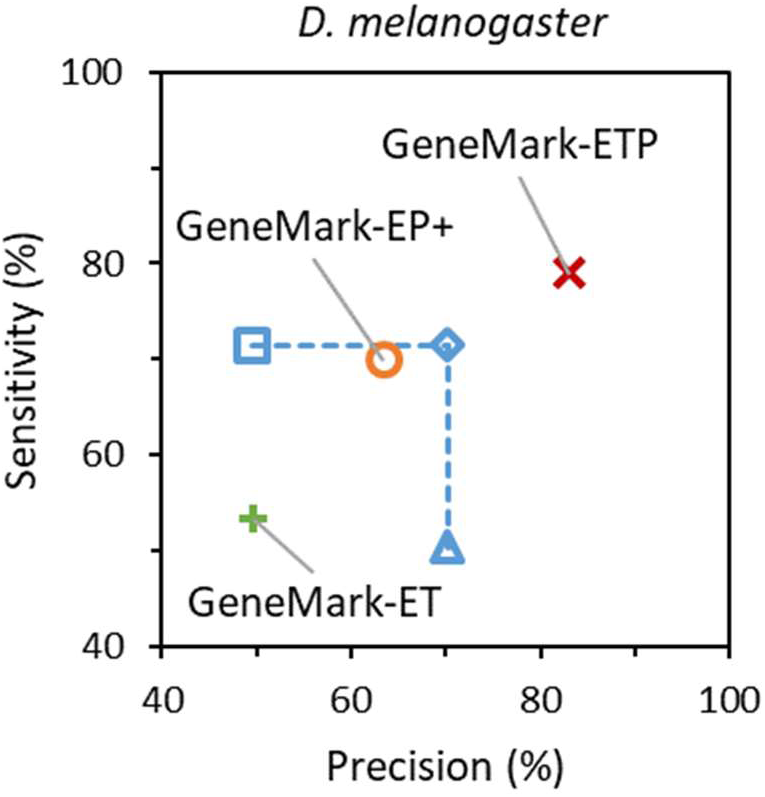
Gene-level Sn and Pr of the gene predictions in the *D. melanogaster* genome. The gene predictions were generated by GeneMark-ET (green), GeneMark-EP+ (orange), and GeneMark-ETP (red). We also show the Sn and Pr values for the Union (square) and the Intersection (triangle) of the sets of GeneMark-ET and GeneMark-EP+ gene predictions and the ‘best combination’ of these two gene prediction sets (diamond). The true positives were defined with respect to the annotation of the *D. melanogaster* genome. The ‘Species excluded’ database was used for GeneMark-EP+ and GeneMark-ETP runs.

Next, the comparisons were made with TSEBRA (Gabriel et al. 2021) and MAKER2 (Holt and Yandell 2011). TSEBRA combines predictions made independently by BRAKER1 and BRAKER2 and filters out less reliable predictions from the union of the sets of genes predicted by the two pipelines. The gene level F1 values of GeneMark-ETP and TSEBRA were comparable in the group of compact genomes. However, the average F1 values of GeneMark-ETP were better by 8.2 points for the two large GC-homogeneous genomes, and by 39.0 points for the two large GC-inhomogeneous genomes, respectively (Figs. 3-4, Supplemental Table S4).

The MAKER2 pipeline runs AUGUSTUS, SNAP, and GeneMark-ES. As extrinsic evidence, MAKER2 uses transcript and protein data. The experiments done with the genomes of *D. melanogaster, D. rerio*, and *M. musculus* representing compact, large GC homogeneous and GC inhomogeneous genomes, demonstrated, respectively, 23.3, 21.7, and 27.8 points improvements in gene level F1, made by GeneMark-ETP (Supplemental Table S5). All the gene finders employed in MAKER2 were run with parameters that, arguably, corresponded to the best-case scenario of training (see Methods). We used specially designed reduced size databases of the cross-species proteins as the runtime of MAKER2 with our minimal size ‘Order excluded’ databases turned out to be prohibitive (see Section S5 of Supplemental Materials). Change to the smaller databases brought F1 of GeneMark-ETP down in comparison with its run with ‘Order excluded’ databases by about 5.0 points for *D. melanogaster* and *M. musculus* but did not change F1 score for *D. rerio. Applicability of the BUSCO scores for assessment of the gene prediction accuracy*

To address possible questions about using the BUSCO scores (Manni et al. 2021) for assessment of gene prediction accuracy, we computed the BUSCO completeness scores both for GeneMark-ETP predictions and for the RefSeq annotations (Supplemental Table S6). The BUSCO scores have reached above 90% for all seven genomes, while the gene level Sensitivity, Precision and the F1 scores, have shown much broader distributions of scores and much larger deviations from the perfect 100 sealing (Figs. 3-4, and Supplemental Tables S3-S4). These results are not unexpected since the BUSCO technique does not expect ‘identity’ between the predicted proteins and the BUSCO proteins that represent a subset of all the proteins in a species proteome. The rules we use in the assessment of the accuracy of gene prediction are more stringent; they require exact matches of predicted and annotated CDSs. While BUSCO scores are overly optimistic if used for assessment of gene prediction quality, they are certainly helpful for the coarse evaluation of completeness of genome assembly and annotation (Huang and Li 2023).

## Discussion

### The GeneMark-ETP features

The GeneMark-ETP algorithm is using several previously developed concepts and approaches, such as the construction of anchored exon borders (GeneMark-ET), generation of external hints from multiple protein spliced alignments (GeneMark-EP+), unsupervised iterative training of GHMM (GeneMark-ES, GeneMark-ET, GeneMark-EP+). However, a distinct feature of GeneMark-ETP is the integration of transcript and protein information in the GeneMarkS-TP module that generates a set of high confidence CDS predictions. The predicted HC CDSs are not changed in the subsequent steps of the algorithm when it makes iterative training and prediction of the CDSs in genomic regions between the HC genes.

We demonstrated that the predicted HC genes had much higher Precision than the initial set of gene candidates (Table 1, Supplemental Fig. S7). The number of HC gene predictions made in each genome with the ‘Order excluded’ database did not differ significantly (except *C. elegans*) from the number of HC genes predicted with the ‘Species excluded’ database (Table 1, Supplemental Fig. S7). Also, the Precision values did not change significantly upon transition from the ‘Order excluded’ to the larger, ‘Species excluded’ databases. These observations indicated that more distant proteins carry significant information for refinements of the HC gene candidates. If for a novel genome there is a limited number of sequenced and annotated genomes of closely related species, then the accuracy estimates made for gene predictions with the ‘Order excluded’ databases would be more realistic than the estimates obtained with the ‘Species excluded’ databases.

Gene prediction Sensitivity also changed in transition from the HC gene candidates to the HC genes. The pattern of change depended on the size of the protein database. For instance, if the ‘Order excluded’ databases were used, then among all the seven genomes only the HC genes predicted in the *C. elegans,* or *D. melanogaster* genomes had lower Sensitivity in comparison with the ones of the HC gene candidates. However, the HC genes predicted with the larger ‘Species excluded’ database, had higher Sensitivity than the HC gene candidates in all the seven genomes (Table 1).

Among groups of gene candidates segregated by the level of extrinsic support, we observed much higher Precision in the group having full extrinsic support than in the group having partial extrinsic support (Table 2). In the three compact genomes and in the tomato genome the magnitude of the difference in Precision was 30-40 percentage points, while in the genomes of *D. rerio, G. gallus,* and *M. musculus* the difference in Precision was more than 50 percentage points. The Precision values in the second and the third groups of gene candidates decreased as the size of the protein database increased (Table 2). This change is explained by the move of many gene candidates from these two groups to the group with full extrinsic support, when the ‘Order excluded’ database was changed to the ‘Species excluded’ one.

Another new feature was application of the two GHMMs i/ the transcript level model used for gene prediction in assembled transcripts and ii/ the genomic level one used for gene prediction in genomic DNA. The training of the transcript level GHMM did follow the path described for GeneMarkS-T (Tang et al. 2015). However, the training process of the genomic GHMM was organized differently in comparison with GeneMark-ES, -ET, -EP+. For those three tools, the initial values of parameters of genomic GHMM were defined by the functions approximating dependence of the *k*-mer frequencies on genomic GC content (Lomsadze et al. 2005; Lomsadze et al. 2014; Bruna et al. 2020). In GeneMark-ETP the training of genomic GHMM is preceded by identification of the HC genes by GeneMarkS-TP (see Methods). Thus, the sequences of the loci containing the HC genes are used as an initial training set for the genomic GHMM with subsequent iterations over the rounds of gene predictions in the non-HC segments and GHMM parameter re-estimation (see Methods, Supplemental Fig. S5). In our experience, the genomic GHHM model derived from the set of HC genes would not change significantly in further iterations, if more than 4,000 HC genes were identified by GeneMarkS-TP.

The genomic GHMM training implemented in GeneMark-ES, -ET, -EP+, used step by step unfreezing of the subsets of the GHMM parameters over the iterations. For instance, the transition probabilities between hidden states, i.e., intron, exon, etc., as well as distributions of durations of hidden states, were fixed during the initial iterations while the values of emission probabilities, derived from the *k*-mer frequencies, were free to change. In the last iterations all the parameters were made free. Such a gradual unfreezing of the parameter set was shown to be unnecessary for GeneMark-ETP, all the GHMM parameters were estimated already in the first iteration.

### Comparisons with other gene finders

As expected, in all the tests GeneMark-ETP performed better than the tools using only a single type of extrinsic evidence, GeneMark-ET, GeneMark-EP+, BRAKER1 and BRAKER2 (see Results, Figs. 3-4). One could notice that GeneMark-EP+, even running with the smaller ‘Order excluded’ database, outperformed GeneMark-ET. This observation was in part due to the fact that GeneMark-ET uses RNA-seq derived hints only in training but not in prediction step. On the other hand, GeneMark-EP+ has a mechanism for enforcing high scoring protein derived external hints into gene predictions made by the Viterbi algorithm.

The magnitudes of improvement that GeneMark-ETP shows in the gene level F1 scores in comparison with the best F1 among GeneMark-ET and GeneMark-EP+ follow already cited patterns of increase from shorter to longer genomes (Figs. 3-4, Supplemental Table S3). For compact genomes, F1 was improved by 11-19 points, and for large genomes – by 26-60 points, with the maximum for the *G. gallus* genome. The gaps in F1 values between GeneMark-ETP and GeneMark-EP+ are smaller when the larger ‘Species excluded’ database was used in comparison with the gaps observed upon the use of the ‘Order excluded’ database (Supplemental Table S3). Certainly, the new features implemented in GeneMark-ETP, as discussed in the previous section, were significant additional factors behind the better performance of GeneMark-ETP in comparison with GeneMark-ET, -EP+, BRAKER1 and BRAKER2 especially for the four large genomes (Figs. 3-4, Supplemental Table S4).

The TSEBRA pipeline selects a subset of all predictions made by either BRAKER1 or BRAKER2 by the rules that attempt to increase Precision without compromising Sensitivity (Gabriel et al. 2021). For compact genomes, *A. thaliana, C. elegance* and *D. melanogaster* TSEBRA shows comparable performance with GeneMark-ETP, with TSEBRA having higher gene level F1 for the first two species (by 1.5 and 4 points) and lower F1 (by 0.5 points) for the last one. However, for the larger genomes: *S. lycopersicum*, *D. rerio*, *G. gallus,* and *M. musculus* GeneMark-ETP improves F1 over TSEBRA by 6.4, 9.0, 45.6, and 29.4 points, respectively (Supplemental Table S4). The double digits of improvements in F1 are observed for the GC inhomogeneous genomes *G. gallus,* and *M. musculus*. All the above-mentioned novel features of GeneMark-ETP contributed to the improvement of the prediction accuracy, especially in the large genomes. Additionally, it should be stated that for large GC-inhomogeneous genome, BRAKER1 and BRAKER2 use a single species-specific GHMM trained in a fashion that effectively makes GHMM for an average genome-specific GC content. GeneMark-ETP constructs three GC specific models for each GC inhomogeneous genome (see Methods). Also, GeneMark-ETP integrates the extrinsic RNA and protein information upon individual prediction of each gene, while in BRAKER1 and BRAKER2 these two channels of extrinsic information are disjointed.

The same novel features of GeneMark-ETP are behind the double-digit improvements in gene level F1 over MAKER2 (Supplemental Table S5). Besides, a number of tools included into the MAKER2 pipeline have been improved since 2011 when MAKER2 was released. Thus, an update of the MAKER2 architecture by transition to more recently developed components would be quite useful.

## Methods

### Data Sets

For computational experiments we selected genomes of seven eukaryotic species. Among them were three model organisms *A. thaliana*, *C. elegans,* and *D. melanogaster,* having well-studied GC-homogeneous and compact in size genomes. The larger genomes of *S. lycopersicum*, and *D. rerio* were GC-homogenous, while the other large genomes of *G. gallus* and *M. musculus* were GC-inhomogeneous. In all cases, sequences of organelles, as well as contigs without chromosome assignment, were excluded (Table 4, Supplemental Table S7).

**Table 4.** The table shows total numbers of protein-coding genes as well as individual CDSs (including alternative isoforms) annotated in the seven genomes (the annotation sources are described in Supplemental Table S7). The numbers of genes and individual CDSs in the intersections of the NCBI RefSeq and the Ensembl annotations are given in parentheses (see Methods). Annotations of *C. elegans, A. thaliana,* and *D. melanogaster* genomes are identical between RefSeq and Ensemble, therefore the intersection sets have the same numbers of genes and CDSs as in the RefSeq annotation.

| Species | Genome length (Mb) | Reference annotation statistics |  |  |  |  |  |
| --- | --- | --- | --- | --- | --- | --- | --- |
|  |  | # of protein-coding genes |  | # of CDSs |  | introns per gene |  |
| *C. elegans* | 100 | 19,969 | - | 28,544 | - | 4.8 | - |
| *A. thaliana* | 119 | 27,445 | - | 40,827 | - | 4.0 | - |
| *D. melanogaster* | 138 | 13,951 | - | 22,395 | - | 2.8 | - |
| *S. lycopersicum* | 807 | 25,158 | (15,138) | 31,911 | (15,150) | 4.4 | (4.3) |
| *D. rerio* | 1,345 | 25,610 | (17,893) | 42,929 | (19,975) | 8.4 | (8.4) |
| *G. gallus* | 1,050 | 17,279 | (10,736) | 38,534 | (12,733) | 9.0 | (9.2) |
| *M. musculus* | 2,723 | 22,405 | (16,531) | 58,318 | (20,708) | 6.0 | (8.6) |

To generate the reference sets of proteins used as a source of extrinsic evidence, we used the OrthoDB v10.1 protein database (Kriventseva et al. 2019). For each of the seven species, we built an initial protein database (PD^0^) containing proteins from the Kingdom, the Phylum or the Class segment of OrthoDB, where the given species belongs (Supplemental Table S8). For each species, we created two reference databases by removing from PD^0^ either i/ proteins of the *species* in question and its strains, the database called ‘Species excluded’, or ii/ proteins of all the species from the same taxonomic *order*, the database called ‘Order excluded’ (see also Bruna et al. 2020). These ‘Order excluded’ and ‘Species excluded’ databases were supposed to mimic practical scenarios when a species of interest would appear at either a *larger* or a *smaller* evolutionary distance from the species present in the protein database. The numbers of proteins in the databases created in our study ranged from 2.6 to 8.3 million (Supplemental Table S8).

Datasets of RNA sequences, the sets of Illumina paired short reads, were selected from the NCBI SRA database. The read length varied between 75 to 151 nt. The total volume of the selected RNA-seq collections varied from 9 Gb for *D. melanogaster* to 83 Gb for *M. musculus* (Supplemental Table S9).

### Algorithm Overview

In the earlier developed GeneMark-ES, -ET, -EP+, estimation of the parameters of GHHMs was done by iterative unsupervised training (Lomsadze et al. 2005; Lomsadze et al. 2014; Bruna et al. 2020). The final set of parameters was used in the final round of gene prediction. The model training and gene prediction are organized differently in GeneMark-ETP which is also an automatic self-training tool (Fig. 1 and Supplemental Methods).

### GeneMarkS-TP: generation of high confidence gene predictions (HC genes)

The wealth of current RNA and protein sequence information precipitates development of a new algorithmic component of a whole genome gene finder, the component conducting gene prediction in assembled transcripts with cross-species protein support. This component of GeneMark-ETP named GeneMarkS-TP is constructed as a pipeline built from several tools. HISAT2 (Kim et al. 2019) splice-aligns the short reads from the selected RNA-seq libraries to the species’ genome. StringTie2 (Kovaka et al. 2019) uses this data for transcript assembly. After filtering out the low-abundance transcripts, the assembled transcripts are merged by StringTie2 into a non-redundant set.

#### Initial gene predictions in transcripts and genomic DNA

A transcript may contain a protein-coding region (CDS) along with 5’ and 3’ UTRs. Predictions of the borders of CDS sequence in a transcript are made by self-training GeneMarkS-T (Tang et al. 2015). Converting the predicting CDS sequence into a chain of protein-coding exons in the genomic DNA, *genomic CDS*, is efficiently done by utilizing data on RNA-seq reads *anchored* to the genome. This information is generated by StringTie2 upon the transcript’s assembly. By combining the coordinates of CDS identified in a transcript with the genomic coordinates of introns produced by StringTie2, the genomic CDS is immediately inferred. The values of Sensitivity and Precision for the predicted genomic CDSs inferred from predictions of GeneMarkS-T in transcripts are shown in Table 1.

#### Refinement of gene predictions

The GeneMarkS-T predictions of *complete* CDSs in the RNA transcripts were shown to be quite accurate (Tang et al. 2015). On the other hand, predictions of *5’ incomplete* CDSs (5’ partial genes) could be less precise. The 5’ partial gene predictions are refined in the GeneMarkS-TP module using protein data.

A 5’ partial CDS is supposed to start from the first nucleotide of a transcript. However, a true complete CDS may reside inside this 5’ partial CDS. To distinguish between the two possibilities, GeneMarkS-TP proceed as follows. Translations of the predicted 5’ partial CDS as well as the longest complete ORF having the same stop of translation as 5’ partial CDS serve as queries in the similarity search by DIAMOND (Buchfink et al. 2015). The protein target common for both searches (E-value<10^-3^) is aligned to both queries. The strengths of sequence conservation along the pairwise alignments are analyzed (Condition S1 in Supplemental Methods). The 5’ partial CDS is confirmed if condition S1 is fulfilled. Otherwise, the predicted 5’ partial CDS is replaced by the shorter complete CDS (see Section S1.1 of Supplemental Methods, Supplemental Tables S11).

#### Selecting HC CDS candidates: CDSs with uniform protein similarity support

Complete and partial CDSs with uniform protein similarity support are selected from the set of CDSs predicted in assembled transcripts. A *complete CDS* is said to have *uniform protein support* if a pairwise alignment of the predicted protein to a known protein satisfies Condition S2 (see Supplemental Methods). A complete CDS with uniform protein similarity support is a candidate for *complete HC CDS*.

One more candidate may be present in the same transcript if it is possible to extend the predicted CDS to the ‘longest’ ORF. If the protein product of this longest ORF has a uniform protein similarity support, (Condition S2, Supplemental Fig. S9), then the extended CDS is yet another candidate for *complete HC CDS,* likely to be an alternative isoform of the same gene.

The uniform protein similarity support could also be found for a predicted *5’ partial CDSs.* If a condition like Condition S2 is fulfilled for a pairwise alignment of the C-terminal of the protein translation of the 5’ partial CDS with a database protein (Supplemental Figs. S10-S11) then the 5’ partial CDS becomes a candidate for a *partial HC CDS*.

#### Selecting HC CDSs candidates: CDSs without uniform protein similarity support

Some genes that do not have uniform protein similarity support may still be qualified as HC CDS candidates. If a predicted *complete* CDS satisfies all the following conditions: (i) the length of the CDS is longer than 299 *nt*, (ii) the 5’ UTR contains in-frame stop codon triplet, and (iii) the exons mapped to genomic DNA do not create a conflict with the ProtHint hints (see Section 2 of Supplemental Methods) then the CDS is designated as a *candidate HC CDS*. Notably, if a CDS lacks a stop codon, it cannot be an HC CDS candidate.

#### Selection of HC CDSs and HC genes from the candidates; generating GeneMarkS-TP output

Alternative transcripts of individual genes are identified by StringTie2 that segregates the whole set of assembled transcripts, complete and incomplete, into groups associated with individual protein-coding genes. We remind that a protein-coding gene is defined here as a set of alternative CDSs coming from the same locus. Alternative mRNA transcripts of a gene may encode identical proteins and have differences only in UTRs. Such alternative transcripts are not considered.

If a gene has only one mRNA transcript (no alternatives), then the HC CDS candidate becomes HC CDS, and the gene is called HC gene. If more than one alternative transcript with HC CDS candidate exist, then we follow the procedure described in Section S3 of Supplemental Materials to select HC CDSs from the candidates and the gene in question becomes the HC gene with thus selected set of HC CDS isoforms. Finally, the set of HC genes that may consist of several alternative HC CDSs, either complete or partial, represents the output of GeneMarkS-TP. This set of HC genes was shown to have significantly higher Precision than the set of initial gene predictions made by GeneMarkS-T (Supplemental Table S11).

### The genomic GHMM training

#### Single step GHMM training

An initial training set for the genomic GHMM is constructed from the set of predicted HC genes. This set, in a GC-inhomogeneous genome, may have significant variation in GC content. To characterize a genome as GC homogenous we check if there exists a 9% wide interval of the GC content distribution that would contain at least 70% of the HC genes selected for training. If there is no such interval, which is a sign that the genomic GC content distribution is wide, the genome is characterized as GC inhomogeneous (Supplemental Fig. S12).

In a *GC homogeneous* genome, the sequences of the predicted HC genes are extended by 1,000nt in both 5’ and 3’ directions (making a set of the *HC loci*). If an HC gene has alternative isoforms, the one with the longest CDS is selected for the training set. The extended sequences including introns and segments of intergenic regions, are used for initial estimation of the genomic GHMM parameters. In a *GC inhomogeneous* genome, the sequences of the *HC loci* are split into three bins: low GC, mid GC, and high GC. The mid GC bin (with the default width of 9%, see Supplemental Fig. S12) is defined by the 9% GC interval that contains the largest number of the HC loci sequences selected for training. Setting up the mid GC interval immediately determines the borders of the low and high GC bins. The sets of the HC loci assigned to the three bins are used to train the GC-specific GHMMs. Note that GeneMarkS-T was developed for gene prediction in GC inhomogeneous transcriptomes. This tool is using a set of GC-specific models (Tang et al. 2015).

#### Extended GHMM training for a genome with homogeneous GC

The logic of extended training of the genomic GHMM is similar but not identical to iterative training used in GeneMark-ET, and -EP+ (Lomsadze et al. 2014; Bruna et al. 2020).

In a *GC homogeneous* genome, the initial GHMM parameters derived from the sequences of the HC loci are used for the first round of gene predictions. The predictions made in the genomic sequences situated between the HC genes, *the non-HC segments*, are of special interest (Supplemental Figs. S5-S6). These predictions could be refined in subsequent iterations, while the predicted HC genes would not change.

The gene predictions made in the non-HC segments are used in ProtHint to generate protein-based hints, as in GeneMark-EP+ (Bruna et al. 2020). An additional set of hints comes from RNA-seq reads mapped to genome by HISAT2 (Kim et al. 2019). All over, there are the following categories of hints: (i) RNA-seq and ProtHint-derived hints to intron borders that agree with each other; (ii) high score ProtHint hints to intron borders; (iii) RNA-seq-based hints to intron borders that may or may not coincide with the intron borders predicted *ab initio*; (iv) hints representing *partial HC genes* that could be extended into the non-HC segments. Note that partial HC genes can appear only at the borders of non-HC segments and technically we consider partial HC genes to belong to non-HC segments.

The hints of the first three categories point to elements of a multi-exon gene without indication that these elements may or may not belong to the same gene. Hints of the fourth category represent ‘chains’ of introns that should belong to the same gene. The requirement of corroboration of RNA-seq hints of category (iii) by *ab initio* predictions allows to filter out the ‘false positive’ intron hints mapped from expressed non-coding RNA. The whole set of selected hints is now ready for enforcement in a run of GeneMark.hmm. The hints are *enforced* in the gene predictions made in the non-HC segments. The predicted genes are included in the updated training set. The iterations stop when the identity of the training sets in two consecutive iterations reaches 99%. The number of iterations observed in our experiments was rarely more than three. Frequently, the convergence has been reached in the second iteration. If the number of HC genes is large (more than 4,000), then, based on our experience, a single iteration is sufficient for convergence. The set of genes predicted in the non-HC segments along with the set of the HC genes made the *final set of genes* predicted by GeneMark-ETP.

#### Extended GHMM training for a genome with inhomogeneous GC

The extended GHMM training for *GC inhomogeneous* genomes works as follows. The three initial GC specific GHMMs are trained on the sequences of the HC loci from the corresponding bins. These GHMMs are used to predict genes in the non-HC segments associated with the same GC bin as the GC specific model. From this point on, the extended training of the three GC-specific GHHM is done on the non-HC segments segregated into the three bins. These trainings are conducted separately following the logic described for the GC homogeneous case. The GC-specific GHMMs are used in the rounds of iterative gene prediction and parameters estimation that occur until convergence (Supplemental Fig. S6). If the number of the HC genes in a bin is large (more than 4,000), then, a single iteration is sufficient for parameter estimation of the GHMM associated with this bin.

In general, GeneMark-ETP could be run in the ‘GC-inhomogeneous’ mode on any genome. However, for a ‘true’ GC-homogeneous genome, this choice would increase runtime and sometimes even decrease gene prediction accuracy due to splitting the overall training set into smaller subsets. Therefore, the degree of GC-heterogeneity is assessed prior to the GHMM training.

### Processing of repetitive elements

Transposable elements (TEs), particularly families of retrotransposons with thousands of copies of similar TE sequences, occupy substantial portions of eukaryotic genomes. Errors in gene prediction may be caused by the presence of repetitive elements with composition similar to protein-coding regions (Yandell and Ence 2012; Torresen et al. 2019). Identification of the repetitive sequences is done independently from gene finding. We generate species-specific repeat libraries *de novo* using RepeatModeler2 (Flynn et al. 2020). Repetitive sequences are then identified in genomic sequence by RepeatMasker (www.repeatmasker.org). Some of the predicted repeats may overlap with protein-coding genes (Bayer et al. 2018). To conduct gene finding in genomic sequence with ‘soft masked’ repeats, a bonus function with species-independent parameter was introduced (Stanke et al. 2008). For a sequence designated as repetitive by a repeat finder, this function increased the likelihood of emission from non-coding state. Here, we have introduced a species-specific penalty function to decrease the likelihood of sequence emission from protein-coding state inside a region identified as a repetitive one (1). The parameter of this function, *q*, is defined (trained) for each genome. Technically, we introduce a state for an overlap of repeat and CDS and compute the probability of a sequence (of length *n*) to be emitted from such a state:

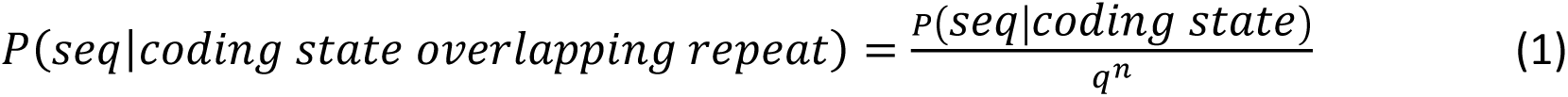

The genome-specific parameter *q* is estimated after obtaining the set of HC genes in the first round of the GHMM training (see Supplemental Methods, Supplemental Fig. S13, Supplemental Table S11).

### Computation of accuracy measures

Assessment of the gene prediction accuracy in each genome was done as follows. The test set for computing the Sensitivity values was the set of genes having identical annotations in NCBI RefSeq and in Ensembl, the intersection set (Table 4). To compute the Precision values, we used the union of genes annotated by NCBI RefSeq and Ensembl. In all the tests, regions of annotated pseudogenes were excluded from consideration. Since annotations of *A. thaliana, C. elegans, and D. melanogaster* genomes were the same at NCBI and Ensemble, then for each of them the union and the intersection sets coincided with each other and with the full complement of annotated genes.

#### GeneMark-ET, GeneMark-EP+, and combinations of their sets of gene prediction

Gene finders, GeneMark-ET or GeneMark-EP+, were designed to use a single source of extrinsic support, either RNA-seq reads or protein database. They were run to get reference points on what accuracy could be achieved with a single source of evidence. We also considered combinations of the sets of genes predicted separately by GeneMark-ET and GeneMark-EP+. The union of the two sets of gene predictions would have increased Sn and decreased Pr. The intersection of the two sets would have increased Pr and decreased Sn (Fig. 5, the case of *D. melanogaster*). The ‘best’ combination of the members of the two sets could be made by removing false positives from the union set or adding true positives made by either method to the intersection set. When the union set is changed by taking away the incorrect predictions, the point for *Union* in Fig. 5 moves horizontally to the right. If one adds correct predictions made by one of the tools but not the other to the intersection set, the *Intersection* point in Fig. 5 moves up vertically. The crossing of the two lines corresponds to the ‘best’ combination of the members of the two sets of predicted genes in terms of (Sn+Pr)/2 (Fig. 5). The described changes cannot be made when a gene finder is running on a novel genome since information on true and false positives is not immediately available. Nevertheless, such modifications could be made for gene predictions in genomes with known annotations.

### Running BRAKER1, BRAKER2, TSEBRA, and MAKER2

To make comparisons with the transcript-supported BRAKER1 (Hoff et al. 2016) and protein-supported BRAKER2 (Bruna et al. 2021) we ran BRAKER1 and BRAKER2 with the same RNA-seq libraries and the same protein databases, respectively, as the ones used in experiments with GeneMark-ETP. Also, we ran TSEBRA (Gabriel et al. 2021), which selects a subset of all the gene predictions made separately by BRAKER1 and BRAKER2 and filters out less reliably predicted genes. TSEBRA was shown to achieve higher accuracy than either BRAKER1 or BRAKER2 running alone.

Execution of MAKER2 has some degree of freedom as the rules of training AUGUSTUS, SNAP and GeneMark.hmm are not precisely specified in the MAKER2 publication (Holt and Yandell 2011). Therefore, we wanted to present the accuracy figures that would, arguably, correspond to the upper limit achieved by the optimal training of MAKER2. The selected models for AUGUSTUS and SNAP were either the models provided by the code developers (available with the respective software distribution), or the models generated by supervised training on the genes annotated by NCBI (RefSeq). As an exception, the SNAP training was done on the Ensembl annotated *D. rerio* genome. Both MAKER2 and GeneMark-ETP use the GeneMark.hmm gene finding algorithm. MAKER2 uses models for GeneMark.hmm self-trained by GeneMark-ES using no extrinsic evidence. We assumed that more precise models would be obtained if extrinsic evidence was also used in the training. Therefore, we trained the GeneMark.hmm models on a set of the HC genes (determined by GeneMark-ETP). To get the accuracy figures, the gene predictions made for the genomes of *D. melanogaster, D. rerio* and *M. musculus* by MAKER2 were processed in the same way as the gene predictions made by GeneMark-ETP or other gene finders. The repeat coordinates and RNA-seq data sets were the same for MAKER2 and GeneMark-ETP (see Supplemental Materials). We compiled reduced size protein databases for running the experiments with MAKER2, since the runtime of MAKER2 sharply increases with the increase in the volume of protein data. The minimal size of the ‘Order Excluded’ database used in the experiments described for the seven genomes was 2.5M proteins. The maximum size database that was used in the experiments for comparison of GeneMark-ETP with MAKER2 was about 300,000 proteins (see Section S5 of the Supplemental Materials). The same reduced in size protein sets were used for both GeneMark-ETP and MAKER2. It should be noted that for a GC-inhomogeneous genome of *M. musculus* GeneMark-ETP used the models with the GC specific parameters. In MAKER2, by design, the GC specific parameters were used in AUGUSTUS but not in SNAP or GeneMark.hmm.

### Runtime of GeneMark-ETP

The runtime of GeneMark-ETP depends linearly on the genome size and is comparable to the one of GeneMark-EP+. The runtime dependence on the size of protein database is also linear. The size of RNA-seq libraries is critical for the HISAT2 runtime but not for the rest of the GeneMark-ETP operations with RNA sequences. We do not include the runtimes of HISAT2 and StringTie2 into the runtime of GeneMark-ETP. To give examples of absolute runtimes, on a machine with 64 CPU cores, GeneMark-ETP finished operations on genomes of *D. melanogaster*, *D. rerio*, and *M. musculus* in 3, 12, and 18 hours, respectively. In these experiments we used the ‘Order excluded’ protein databases and RNA reads sets described in Supplemental Tables S8-S9.

### Software availability

GeneMark-ETP is available at https://exon.gatech.edu/GeneMark/license_download.cgi and in Supplemental Materials. All scripts and data used to generate figures and tables in this manuscript are available at https://github.com/gatech-genemark/GeneMark-ETP-exp and in Supplemental Materials. GeneMark-ETP was included in the recently developed pipeline BRAKER3 (Gabriel et al. 2023).

### Competing Interests Statement

GeneMark-ETP and its part GeneMark.hmm, are distributed under the Creative Commons license. Licensing GeneMark.hmm by commercial companies may create a conflict of interest for AL and MB. TB declares having no competing interests.

## Acknowledgements

The National Institutes of Health [GM128145 to MB] supported this work. Georgia Tech provided funding for the open access charge.

## Author contributions

T.B, A.L. and M.B designed the project; T.B and A.L. designed and wrote the code, prepared all the initial data, and conducted computational experiments; T.B, A.L. and M.B wrote and edited the manuscript.

## Notes

### Summary of Updates

The manuscript was improved for clarity.

## References

Allen JE, Pertea M, Salzberg SL. 2004. Computational gene prediction using multiple sources of evidence. Genome Research 14: 142–148.

Allen JE, Salzberg SL. 2005. JIGSAW: integration of multiple sources of evidence for gene prediction. Bioinformatics 21: 3596–3603.

Banerjee S, Bhandary P, Woodhouse M, Sen TZ, Wise RP, Andorf CM. 2021. FINDER: an automated software package to annotate eukaryotic genes from RNA-Seq data and associated protein sequences. BMC Bioinformatics 22: 205.

Bayer PE, Edwards D, Batley J. 2018. Bias in resistance gene prediction due to repeat masking. Nat Plants 4: 762–765.

Bruna T, Hoff KJ, Lomsadze A, Stanke M, Borodovsky M. 2021. BRAKER2: automatic eukaryotic genome annotation with GeneMark-EP+ and AUGUSTUS supported by a protein database. NAR Genom Bioinform 3: lqaa108.

Bruna T, Lomsadze A, Borodovsky M. 2020. GeneMark-EP+: eukaryotic gene prediction with self-training in the space of genes and proteins. NAR Genom Bioinform 2: lqaa026.

Buchfink B, Xie C, Huson DH. 2015. Fast and sensitive protein alignment using DIAMOND. Nature Methods 12: 59–60.

Burge C, Karlin S. 1997. Prediction of complete gene structures in human genomic DNA. J Mol Biol 268: 78–94.

Coghlan A, Fiedler TJ, McKay SJ, Flicek P, Harris TW, Blasiar D, n GC, Stein LD. 2008. nGASP--the nematode genome annotation assessment project. BMC Bioinformatics 9: 549.

Cook DE, Valle-Inclan JE, Pajoro A, Rovenich H, Thomma B, Faino L. 2019. Long-Read Annotation: Automated Eukaryotic Genome Annotation Based on Long-Read cDNA Sequencing. Plant Physiol 179: 38–54.

Flynn JM, Hubley R, Goubert C, Rosen J, Clark AG, Feschotte C, Smit AF. 2020. RepeatModeler2 for automated genomic discovery of transposable element families. Proc Natl Acad Sci U S A 117: 9451–9457.

Gabriel L, Bruna T, Hoff KJ, Ebel M, Lomsadze A, Borodovsky M, Stanke M. 2023. BRAKER3: Fully Automated Genome Annotation Using RNA-Seq and Protein Evidence with GeneMark-ETP, AUGUSTUS and TSEBRA. bioRxiv doi:10.1101/2023.06.10.544449: 2023.2006.2010.544449.

Gabriel L, Hoff KJ, Bruna T, Borodovsky M, Stanke M. 2021. TSEBRA: transcript selector for BRAKER. Bmc Bioinformatics 22.

Goodswen SJ, Kennedy PJ, Ellis JT. 2012. Evaluating high-throughput ab initio gene finders to discover proteins encoded in eukaryotic pathogen genomes missed by laboratory techniques. PLoS One 7: e50609.

Gremme G, Brendel V, Sparks ME, Kurtz S. 2005. Engineering a software tool for gene structure prediction in higher organisms. Inform Software Tech 47: 965–978.

Guigo R, Flicek P, Abril JF, Reymond A, Lagarde J, Denoeud F, Antonarakis S, Ashburner M, Bajic VB, Birney E et al. 2006. EGASP: the human ENCODE Genome Annotation Assessment Project. Genome Biol 7 Suppl 1: S2 1-31.

Haas BJ, Salzberg SL, Zhu W, Pertea M, Allen JE, Orvis J, White O, Buell CR, Wortman JR. 2008. Automated eukaryotic gene structure annotation using EVidenceModeler and the Program to Assemble Spliced Alignments. Genome Biol 9: R7.

Hoff KJ, Lange S, Lomsadze A, Borodovsky M, Stanke M. 2016. BRAKER1: Unsupervised RNA-Seq-Based Genome Annotation with GeneMark-ET and AUGUSTUS. Bioinformatics 32: 767–769.

Holt C, Yandell M. 2011. MAKER2: an annotation pipeline and genome-database management tool for second-generation genome projects. BMC Bioinformatics 12: 491.

Howe KL, Chothia T, Durbin R. 2002. GAZE: A generic framework for the integration of gene-prediction data by dynamic programming. Genome Research 12: 1418–1427.

Huang N, Li H. 2023. miniBUSCO: a faster and more accurate reimplementation of BUSCO. bioRxivdoi:10.1101/2023.06.03.543588.

Keilwagen J, Hartung F, Paulini M, Twardziok SO, Grau J. 2018. Combining RNA-seq data and homology-based gene prediction for plants, animals and fungi. BMC Bioinformatics 19: 189.

Kim D, Paggi JM, Park C, Bennett C, Salzberg SL. 2019. Graph-based genome alignment and genotyping with HISAT2 and HISAT-genotype. Nat Biotechnol 37: 907–915.

Kiryutin B, Souvorov A, Tatusova T. 2007. Prosplign: protein to genomic alignment tool. In 11th Annual International Conference in Research in Computational Molecular Biology, San Francisco, USA.

Kong J, Huh S, Won JI, Yoon J, Kim B, Kim K. 2019. GAAP: A Genome Assembly + Annotation Pipeline. Biomed Res Int 2019: 4767354.

Kovaka S, Zimin AV, Pertea GM, Razaghi R, Salzberg SL, Pertea M. 2019. Transcriptome assembly from long-read RNA-seq alignments with StringTie2. Genome Biol 20: 278.

Kriventseva EV, Kuznetsov D, Tegenfeldt F, Manni M, Dias R, Simao FA, Zdobnov EM. 2019. OrthoDB v10: sampling the diversity of animal, plant, fungal, protist, bacterial and viral genomes for evolutionary and functional annotations of orthologs. Nucleic Acids Research 47: D807–D811.

Kulp D, Haussler D, Reese MG, Eeckman FH. 1996. A generalized hidden Markov model for the recognition of human genes in DNA. Proc Int Conf Intell Syst Mol Biol 4: 134–142.

Lewin HA, Richards S, Lieberman Aiden E, Allende ML, Archibald JM, Balint M, Barker KB, Baumgartner B, Belov K, Bertorelle G et al. 2022. The Earth BioGenome Project 2020: Starting the clock. Proc Natl Acad Sci U S A 119.

Liu Q, Mackey AJ, Roos DS, Pereira FC. 2008. Evigan: a hidden variable model for integrating gene evidence for eukaryotic gene prediction. Bioinformatics 24: 597–605.

Lomsadze A, Burns PD, Borodovsky M. 2014. Integration of mapped RNA-Seq reads into automatic training of eukaryotic gene finding algorithm. Nucleic Acids Res 42: e119.

Lomsadze A, Ter-Hovhannisyan V, Chernoff YO, Borodovsky M. 2005. Gene identification in novel eukaryotic genomes by self-training algorithm. Nucleic Acids Res 33: 6494–6506.

Manni M, Berkeley MR, Seppey M, Simao FA, Zdobnov EM. 2021. BUSCO Update: Novel and Streamlined Workflows along with Broader and Deeper Phylogenetic Coverage for Scoring of Eukaryotic, Prokaryotic, and Viral Genomes. Mol Biol Evol 38: 4647–4654.

Parra G, Blanco E, Guigo R. 2000. GeneID in Drosophila. Genome Res 10: 511–515.

Pertea M, Pertea GM, Antonescu CM, Chang TC, Mendell JT, Salzberg SL. 2015. StringTie enables improved reconstruction of a transcriptome from RNA-seq reads. Nat Biotechnol 33: 290–295.

Scalzitti N, Jeannin-Girardon A, Collet P, Poch O, Thompson JD. 2020. A benchmark study of ab initio gene prediction methods in diverse eukaryotic organisms. BMC Genomics 21: 293.

Slater GS, Birney E. 2005. Automated generation of heuristics for biological sequence comparison. BMC Bioinformatics 6: 31.

Song L, Sabunciyan S, Yang G, Florea L. 2019. A multi-sample approach increases the accuracy of transcript assembly. Nat Commun 10: 5000.

Stanke M, Diekhans M, Baertsch R, Haussler D. 2008. Using native and syntenically mapped cDNA alignments to improve de novo gene finding. Bioinformatics 24: 637–644.

Tang S, Lomsadze A, Borodovsky M. 2015. Identification of protein coding regions in RNA transcripts. Nucleic Acids Res 43: e78.

Ter-Hovhannisyan V, Lomsadze A, Chernoff YO, Borodovsky M. 2008. Gene prediction in novel fungal genomes using an ab initio algorithm with unsupervised training. Genome Res 18: 1979–1990.

Torresen OK, Star B, Mier P, Andrade-Navarro MA, Bateman A, Jarnot P, Gruca A, Grynberg M, Kajava AV, Promponas VJ et al. 2019. Tandem repeats lead to sequence assembly errors and impose multi-level challenges for genome and protein databases. Nucleic Acids Res 47: 10994–11006.

Trapnell C, Williams BA, Pertea G, Mortazavi A, Kwan G, van Baren MJ, Salzberg SL, Wold BJ, Pachter L. 2010. Transcript assembly and quantification by RNA-Seq reveals unannotated transcripts and isoform switching during cell differentiation. Nat Biotechnol 28: 511–515.

Yandell M, Ence D. 2012. A beginner’s guide to eukaryotic genome annotation. Nat Rev Genet 13: 329–342.

Zickmann F, Renard BY. 2015. IPred - integrating ab initio and evidence based gene predictions to improve prediction accuracy. BMC Genomics 16: 134.

