## supplementary for "A new gene finding tool GeneMark-ETP significantly improves the accuracy of automatic annotation of large eukaryotic genomes"

### Supplemental Material

### Supplemental Methods

#### S1. GeneMarkS-TP: predicting genes in RNA transcripts with protein database support.

##### S1.1 Corrections of the 5' end gene predictions

The CDS prediction in assembled transcripts is done by GeneMarkS-T (Tang et al. 2015). We have observed that GeneMarkS-T made very few errors when predicting 5' complete CDSs, those having start codons within transcripts. On the other hand, the 5' incomplete CDSs predicted by GeneMarkS-T with the start codons residing near the first nucleotide of a transcript carry more frequent errors that should be corrected. We need to discriminate between a correctly predicted 5' incomplete CDS and an incorrect 5' incomplete CDS with a true complete CDS residing inside.

Incomplete CDSs predicted by GeneMarkS-T in transcripts serve as queries in searches for homologous proteins (targets) in a reference protein database (e.g., by DIAMOND (Buchfink et al. 2015)). If among the similarity search hits (targets) exists at least one target that i/ is common for both queries and ii/ shows *better support* for the 5' partial CDS, *the 5' partial CDS is predicted*.

Otherwise, the CDS starting with the internal ATG is selected as the *predicted complete CDS*. If the sets of protein targets in the two searches (those with 25 best scores, the default setting) do not overlap, the 5' partial CDS is selected. If both similarity searches do not produce targets, then the transcript is removed from consideration.

The quantitative meaning of the *better support* provided by the protein alignment data is formalized by the following condition:

$$(b - a) - (a - 1) > 1000 * \ln \frac{AAI_{complete}}{AAI_{partial}} \qquad \text{(S1)}$$

Here, *a* and *b* are the starting positions of the local alignments within the target protein for the longer and shorter protein queries respectively (see Supplemental Fig. S10 and S11). $AAI_{partial}$ and $AAI_{complete}$ are, respectively, the percentages of *amino acid identities* in the alignments of the longer and shorter query proteins with the target protein. $AAI_{partial}$ is defined within the range "a-c", $AAI_{complete}$ is defined within the range "b-c", where c is the common end position of the two local pairwise alignments (Supplemental Figs. S10 and S11).

If condition (S1) is fulfilled, the longer query is selected, the 5' partial CDS.
If condition (S1) is not fulfilled, the shorter query, a *complete* CDS is selected.

Notably, "a-1" is the length of the unaligned N proximal part of the long query.
A large "a-1" is likely to indicate the presence of a translated 5' UTR region situated upstream of a complete gene. A small "b-a" indicates that an extension of the complete gene candidate does not extend the zone of two proteins similarity, again a support for the complete gene prediction.

The larger value of the *AAI* ratio, the more conservation exists between query and target protein subsequences in the range "b-a". Therefore, the increase in the *AAI* ratio favors the 5' partial candidate. The *AAI* ratio is scaled using a logarithm with a factor of 1,000, i.e., 1,000*log(…).

#### S1.2 Removal of the 3' partial CDS predictions

The 3' partial predictions were rarely observed. This low frequency could be expected since RNA-Seq libraries used in our experiments, prepared with the poly-A tail enrichment of mRNA transcripts, should predominantly carry 3' end complete transcripts (Zhao et al. 2014). Therefore, all the 3' partial genes were removed from the list of candidates for high-confidence CDSs.

#### S1.3. Extensions of GeneMarkS-T CDS predictions to the longest ORFs

Most eukaryotic genes are translated from the ATG start codon closest to the transcript 5' end (Kozak 1999). Still, the translation can be initiated at one of the downstream ATG starts, e.g., when the most upstream start has a weak translation initiation signal known as the Kozak pattern (Kozak 1987). GeneMarkS-T computes Kozak pattern score (with respect to the model with parameters derived in species-specific self-training) to account for the possibility of non-5'-most translation start codons. However, the Kozak pattern is relatively weak. We have observed that the predictions of CDSs with non-5'-most start codons carry a higher false-positive rate than the predictions of CDSs with 5'-most start codons. Therefore, we use the following rule. If a CDS predicted in a transcript could be extended to the 5'-most start codon, and the translation of this extension is supported by alignment to a target protein, the extended predicted CDS is considered a candidate for an HC CDS along with the one with non-5'-most start.

#### S1.4 Complete genes with uniform protein support

In the described above similarity searches we have dealt with local pairwise alignments. Still, being interested in the accurate prediction of all protein-coding exons, we are concerned about *uniform* protein support showing evolutionary conservation over the whole protein-coding region. We say that a *uniform protein support* exists for a predicted *complete* CDS if there is a significant BLASTp alignment (with E-value better than $10^{-3}$) of the translation of the predicted CDS $Q$ to a protein in a database $T$ and the following condition is satisfied:

$$(|Q_{start} - T_{start}| \leq 5) \wedge (|(Q_{len} - Q_{end}) - (T_{len} - T_{end})| \leq 20) \qquad \text{(S2)}$$

Here, $Q_{start}, Q_{end}$, $(T_{start}, T_{end})$ are, respectively, the positions of the start and end of the alignment within the query protein (within the target protein); $Q_{len}, T_{len}$ are the lengths of the query and target proteins, respectively (Supplemental Fig. S9).

Experiments with multiple sequence alignments (MSA) of orthologous proteins demonstrated that internal sections of MSA were usually most conserved, while the N-proximal regions of the proteins were less conserved, and the least conserved regions in MSA were usually C-proximal regions. Therefore, testing for conservation of the N- and C- proximal regions provided sufficient evidence of evolutionary conservation across the pair of proteins. Condition S2 allows some misalignment at the alignment start and even to a larger degree at the alignment end. Predicted CDS is called a complete CDS with uniform protein support if a translated query protein has an alignment to at least one target (out of the best scored 25, the default setting) that satisfies condition S2. All such predicted CDSs are included in the set of high-confidence CDSs.

#### S1.5 Tests of conditions S1 and S2

To assess the degree of improvement in the quality of gene sets selected with conditions S1 and S2, we used the following approach. We have prepared test sets of transcript sequences with complete and partial CDSs. The ground-truth labels were determined from reference annotations. GeneMarkS-T was run on these sequences. Next, for each transcript, the alignments of the longer and shorter queries with the target proteins were made, and the features used in conditions S1 and S2 were selected. We assessed the efficiency of the empirical rules for selecting partial and complete CDSs (Condition S1) as well as selecting CDSs with uniform protein support (Condition S2) with the efficiency of two other possible approaches. We trained random forest and logistic regression classifiers (with Python's scikit-learn machine learning library) using all alignment features offered by DIAMOND's tabular output (Buchfink et al. 2015) i/ to classify CDS predictions as complete or partial (compared to the use of condition S1), ii/ to claim uniform protein support (Compared to the use of condition S2). The training sets for the two ML methods did not overlap with the test set. We observed that the use of conditions S1 and S2 produced more accurate results than the results generated by the application of general-purpose random forest or logistic regression models (data not shown).

#### S2. ProtHint filter for high-confidence gene candidates (in the *ab initio* category)

Some GeneMarkS-T predicted CDSs not uniformly supported by proteins (and not satisfying Condition S2) could still be included in the set of HC CDSs. Such predictions should satisfy several conditions (see Main text), one of which is no contradiction to the ProtHint hints. To detect such a conflict, we proceed as follows. First, a CDS predicted by GeneMarkS-T is mapped to genomic DNA. Second, the translation of the initially predicted CDS and its genomic locus is used by ProtHint as the protein and CDS seeds to generate hints for the next round of CDS prediction in the same locus (Bruna et al. 2020). Next, the borders of the thus determined exons are compared to the ProtHint hints. We say that the contradiction exists if (i) at least one of ProtHint's introns overlaps a mapped exon, or (ii) a ProtHint defined stop codon overlaps an exon or intron of the mapped gene, or (iii) a ProtHint start codon overlaps an exon or intron of the mapped gene (except the start-to-start overlap).

#### S3. Alternative HC CDSs

An additional round of selection is made to subject CDSs that satisfy Condition S2 to a more stringent restriction. Let $I^g_{complete}$ be a set of complete alternative CDSs of protein-coding gene $g$ and $I^g_{partial}$ is a set of its alternative partial CDSs. Each isoform $i$ is assigned a score $s(i)$ -- the *bitscore* of its best hit to a protein in the protein database. We compute the maximum score of all the complete CDS isoforms for a gene $g$, denoted as $\mathrm{s}(g_{complete})$. A score of a CDS isoform $\mathrm{s}(i)$ selected as complete HC CDS isoform must satisfy the inequality:

$$\mathrm{s}(i) \geq 0.8 \times \mathrm{s}(g_{complete}) \quad (i \in I^g_{complete}) \qquad \text{(S3)}$$

Among the partial alternative CDSs of gene $g$, we determine the maximum score $\mathrm{s}(g_{partial})$. If $\mathrm{s}(g_{partial})$ is larger than $\mathrm{s}(g_{complete})$, the partial CDS isoform with this largest score is selected as the partial HC isoform. In this case, all the complete HC isoforms are removed. Otherwise, if $\mathrm{s}(g_{partial})$, is lower than $\mathrm{s}(g_{complete})$, then only complete HC CDSs of gene $g$ are retained.

If all alternative HC CDS candidates were defined *ab initio*, then the one with the longest protein-coding region is selected as the predicted HC CDS.

The numbers of predicted alternative CDSs are smaller than the numbers of annotated alternative CDSs (Table 3), because we predict alternative CDSs only for the HC genes, a subset of all genes. Moreover, the CDS isoforms of the HC genes must have full protein support (Condition S3) which further limits the number of predicted CDS isoforms.

#### S4. Computing the species-specific repeat penalty parameter

For each genome, after identification of the HC CDSs and the first iteration of the GHMM model training, we estimate species-specific parameter $q$.

We have the set of the HC CDSs, the first version of the full GHMM model, and the coordinates of the repeats identified in genomic DNA. GeneMark.hmm is run several times with different $q$ values to predict CDSs in the genomic sequences containing the HC CDSs for which we compute the gene level F1 value (Supplemental Fig. S12-A). The value $q$ delivering the F1 maximum was chosen as the species-specific repeat penalty. We have shown that this value is close to $q$ found when the test set of CDSs is made based on genome annotation. We also observed that the value $q$ was robust with respect to the size of the HC CDSs set (data not shown).

Moreover, we have found that the use of the exon level Sn led to more robust estimation of $q$ in comparison with use of the gene level F1 (data not shown). Practically, we first find the $q'$ value maximizing the number of correctly predicted exons in the set of HC genes, $e_{max}$ (Supplemental Fig. S12-B). Then, the value $q*$ at which $0.998 \times e_{max}$ exons are correctly predicted (marked for *A. thaliana* and *D. melanogaster* in panel A of Supplemental Fig. S12-A) is selected as $q$. To reduce the runtime of the repeat penalty parameter estimation, we use simulated annealing (Kirkpatrick et al. 1983).

**S5. Data sets used in computational experiments with MAKER2**

Three model organisms having different types of genome organization were selected:

- *Drosophila melanogaster* – compact, GC homogeneous genome.
- *Danio rerio* – large, GC homogenous genome
- *Mus musculus* – large, GC heterogeneous genome

The following information was available to MAKER2.

Repeat coordinates predicted by RepeatMasker in the MAKER2 supported GFF format:

```
rmasker_out2maker_gff.pl < genome.fasta.out > repeatmasker.gff
```

Transcripts assembled from RNA-Seq by HISAT2/StringTie2 were provided as transcriptome input to MAKER2 (the same input as in the GeneMark-ETP runs)

As a protein database input for both MAKER2 and GeneMark-ETP we used the OrthoDB proteins as follows:

For *Drosophila melanogaster,* 274,283 proteins from

*Drosophila ananassae*
*Drosophila biarmipes*
*Drosophila bipectinate*
*Drosophila busckii*
*Drosophila elegans*
*Drosophila erecta*
*Drosophila eugracilis*
*Drosophila ficusphila*
*Drosophila grimshawi*
*Drosophila hydei*

*Drosophila mojavensis*
*Drosophila obscura*
*Drosophila pseudoobscura*
*Drosophila rhopaloa*
*Drosophila serrata*
*Drosophila takahashii*
*Drosophila virilis*
*Drosophila willistoni*
*Drosophila yakuba*

For *Danio rerio,* 181,842 proteins from:

*Cyprinus carpio*
*Sinocyclocheilus anshuiensis*
*Sinocyclocheilus 6ahari*
*Sinocyclocheilus rhinocerous*

For *Mus musculus,* 207,553 proteins from:

*Cavia porcellus*
*Cricetulus griseus*
*Fukomys damarensis*
*Ictidomys tridecemlineatus*
*Marmota marmota marmota*
*Mesocricetus auratus*
*Mus caroli*
*Mus 6ahari*
*Octodon degus*
*Rattus norvegicus*

MAKER2 was executed with the gene finders AUGUSTUS, GeneMark.hmm and SNAP.
The following model files were used by the gene finders:

For *Drosophila melanogaster*:
AUGUSTUS – "fly" from the AUGUSTUS distribution.
GeneMark.hmm – model created by GeneMark-ETP.
SNAP – "D.melanogaster.hmm" from the SNAP distribution.

For *Danio rerio*:
AUGUSTUS – the "zebrafish" model from the AUGUSTUS distribution.
GeneMark.hmm – the model created by GeneMark-ETP.
SNAP – the model trained according to instructions from the SNAP distribution. The training set matched the test set used for evaluation of the MAKER2 performance.

For *Mus musculus*:
AUGUSTUS – the "human" model from the AUGUSTUS distribution.
GeneMark.hmm – the 'medium GC' model created by GeneMark-ETP for the *Mus musculus* genome.

SNAP – the “mam46.hmm” mammalian model for the medium GC bin from SNAP distribution.

MAKER2 was executed with the following settings in the MAKER2 configuration file:

genome=genome.fasta
est=transcriptome.fasta
protein=proteindb.fasta
model_org=   #empty
rm_gff=repeatmasker.gff
snaphmm=snap.model
gmhmm=genemark.mod
augustus_species=model_name
est2genome=1
protein2genome=1
alt_splice=1
always_complete=1
keep_preds=1 for D. melanogaster
keep_preds=0 for D. rerio and M. musculus
split_hit=20000
max_dna_len=1000000

A LINUX node with 96 cores was used to execute MAKER2 in MPI mode at the Azure cloud.

The gene prediction accuracy of MAKER2 and GeneMark-ETP (Supplemental Table S5) was estimated as described in the main text (see Methods).

### Supplemental Figures

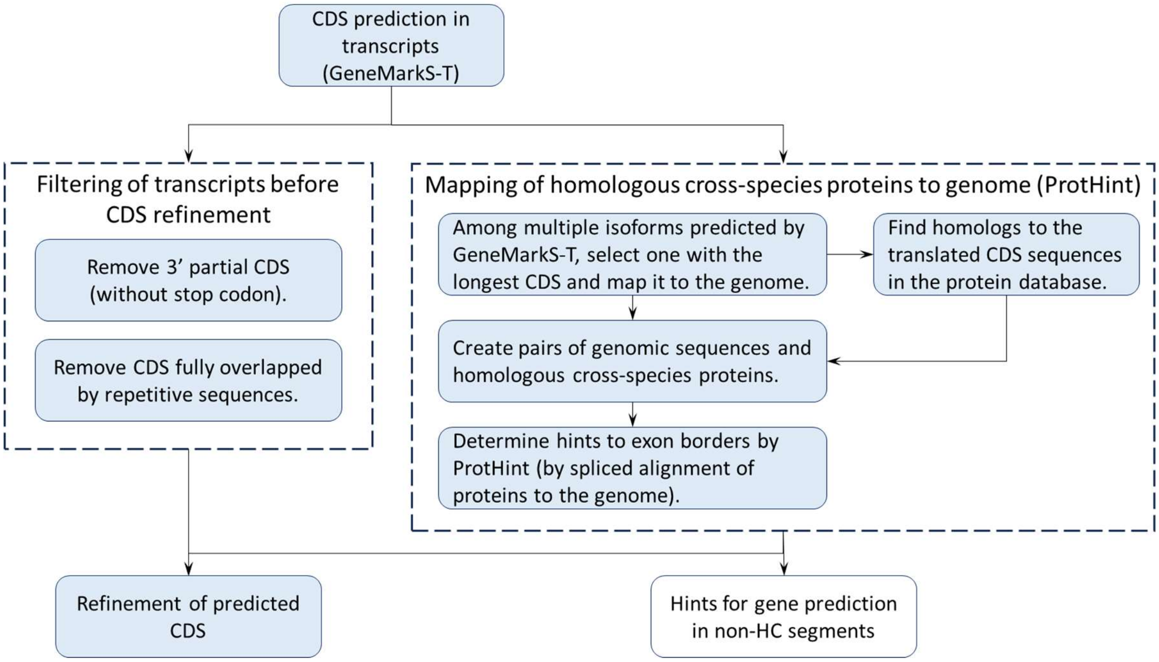


**Supplemental Figure S1.** Schematics showing the transcript processing steps in GeneMarkS-TP (see Fig. 2).

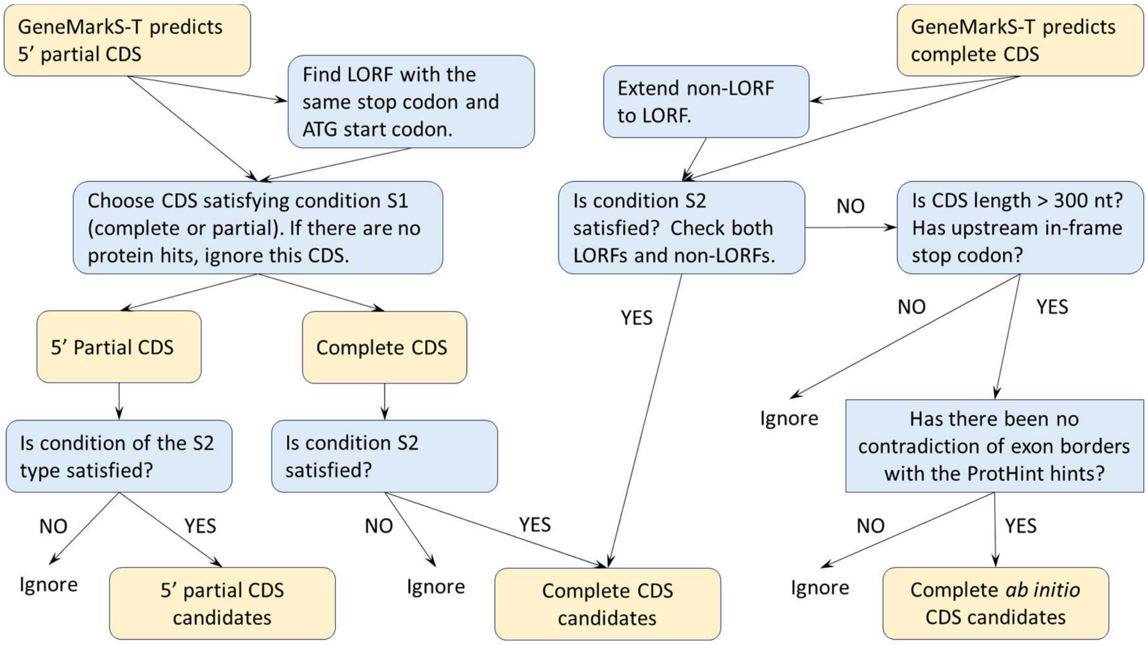


**Supplemental Figure S2.** Schematics of the generation of HC CDS candidates in GeneMarkS-TP (the refinement block in Fig. 2).

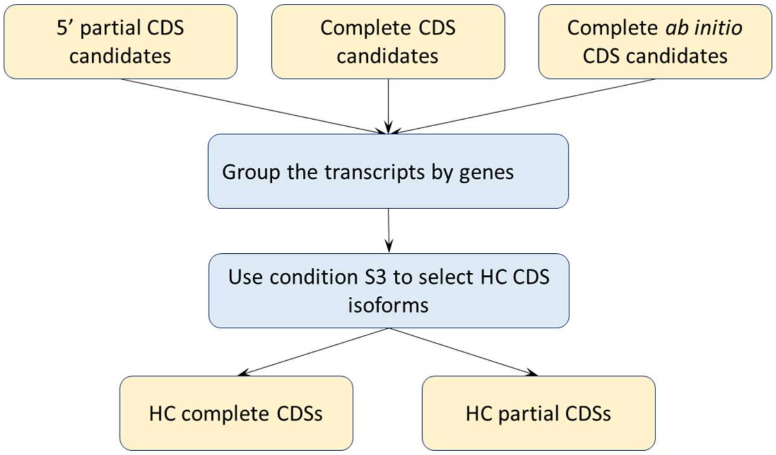


**Supplemental Figure S3.** Schematics of the selection of HC CDSs in GeneMarkS-TP (see Fig.2).

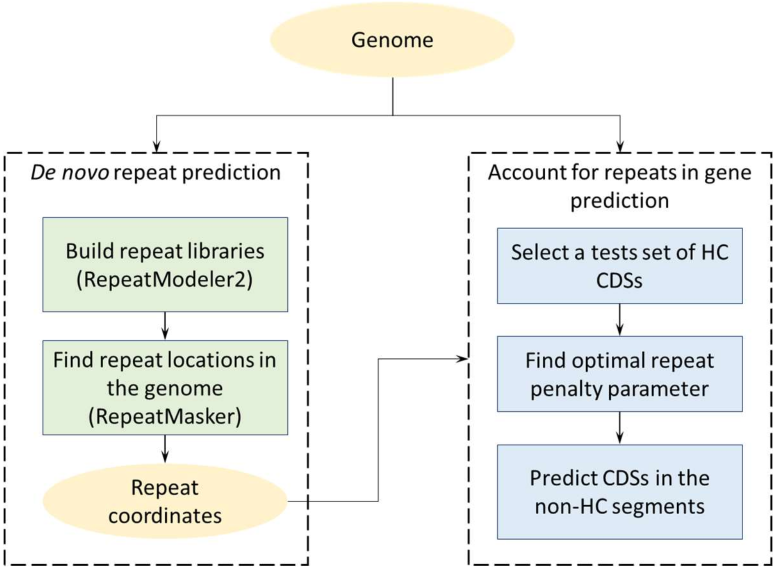


**Supplemental Figure S4.** Schematics of the repetitive sequence identification and processing. *De novo* repeats prediction module (shown on the left) is not a part of GeneMark-ETP (see Fig.1).

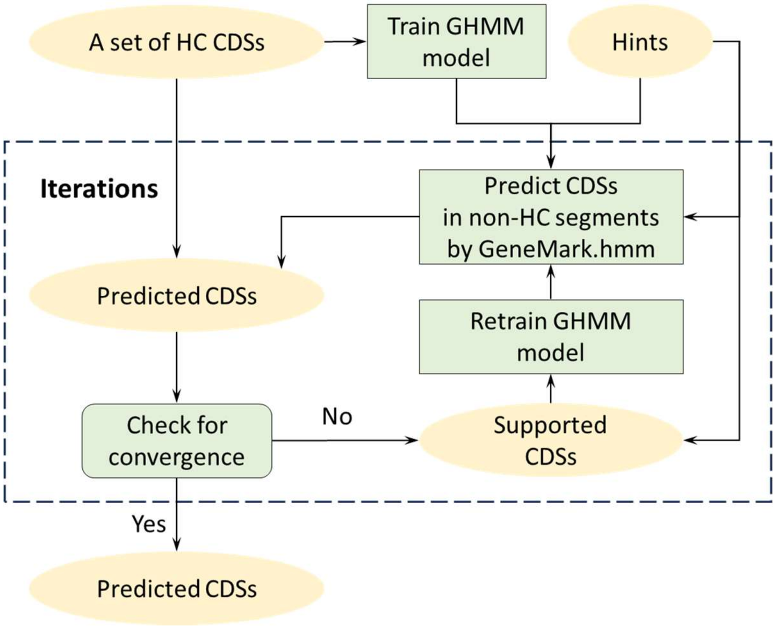


**Supplemental Figure S5.** Workflow of the training of the GHMM model used in GeneMark.hmm (see Fig.1).

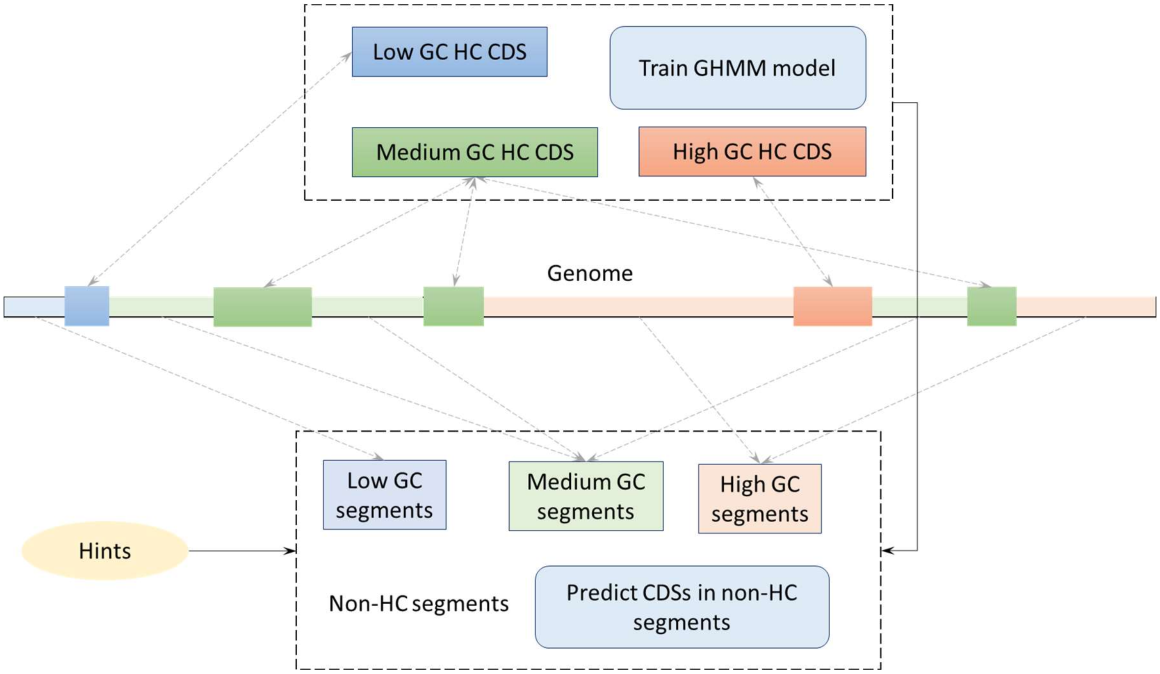


**Supplemental Figure S6.** Schematics of the identification and the use of the non-HC segments in the GHMM training and CDS prediction (see Fig.1).

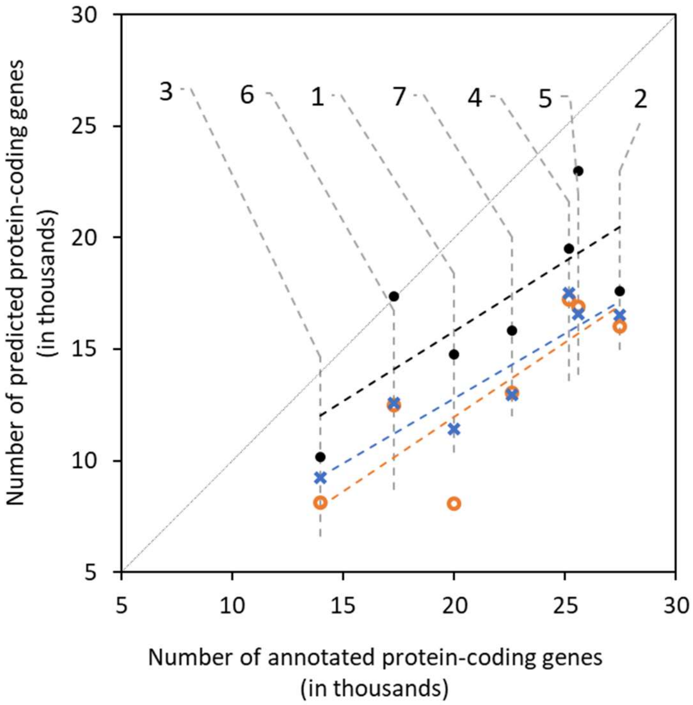


**Supplemental Figure S7.** Numbers of protein-coding genes predicted at initial stages of running GeneMark-ETP (i) genes predicted in assembled transcripts by GeneMarkS-T (black dots), (ii) HC genes predicted by GeneMarkS-TP with the 'Order excluded' protein database (orange circles) and with the 'Species excluded' database (blue crosses). The number of genes annotated in each genome is taken from the RefSeq annotation (Supplemental Table S7). The numerical designations of the species are as follows: 1 - *C. elegans*, 2 - *A. thaliana*, 3 - *D. melanogaster*, 4 - *S. lycopersicum*, 5 - *D. rerio*, 6 - *G. gallus*, 7 - *M. musculus*.

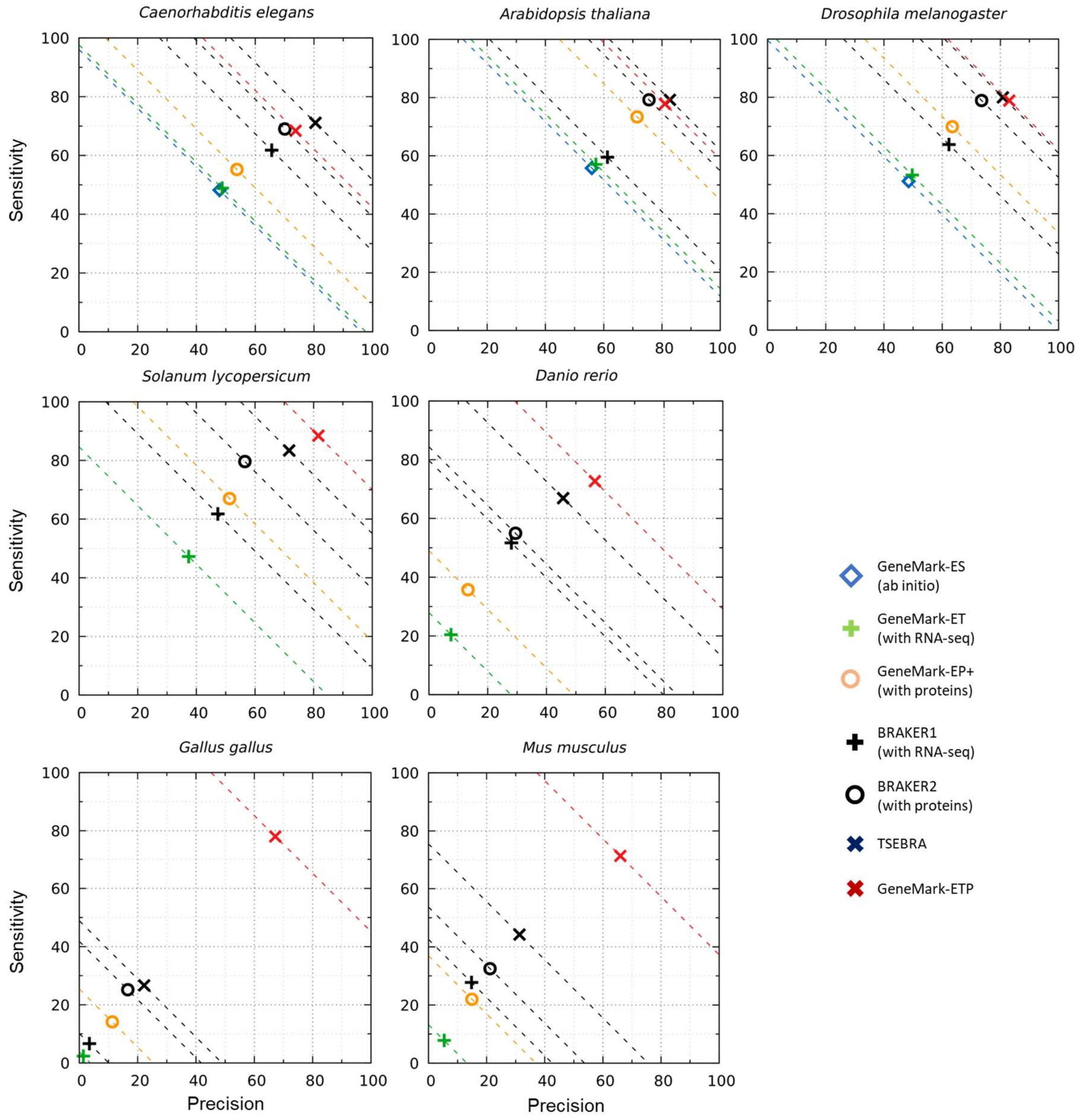


**Supplemental Figure S8.** Gene level accuracy of the seven gene prediction tools (see legends to Figs. 3-4). Compared to the figures in the main text, where we used the 'Order excluded' protein databases for each species, here we used the larger 'Species excluded' databases.

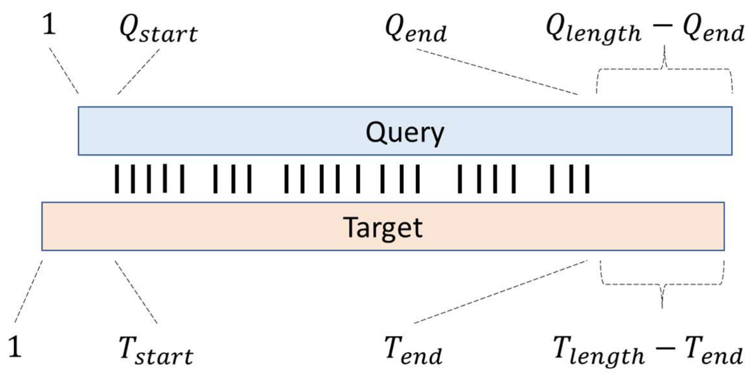


**Supplemental Figure S9**. Depictions of the alignment features used in Condition S2.

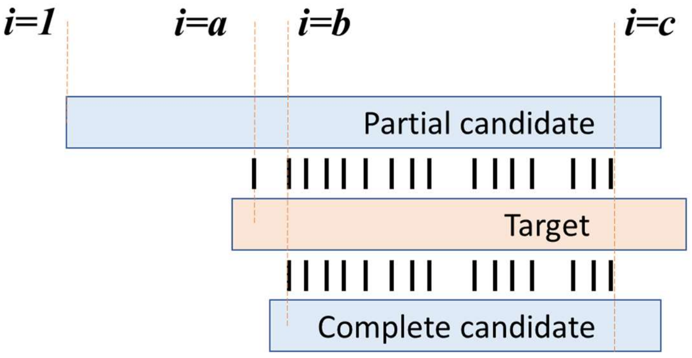


**Supplemental Figure S10.** Illustration for the case when Condition S1 is not fulfilled, and the GeneMarkS-T prediction is classified as *complete CDS*. Here *a* and *b* are positions of the starts of the local alignments of respective longer and shorter protein queries, while *c* is the end position of the local pairwise alignments.

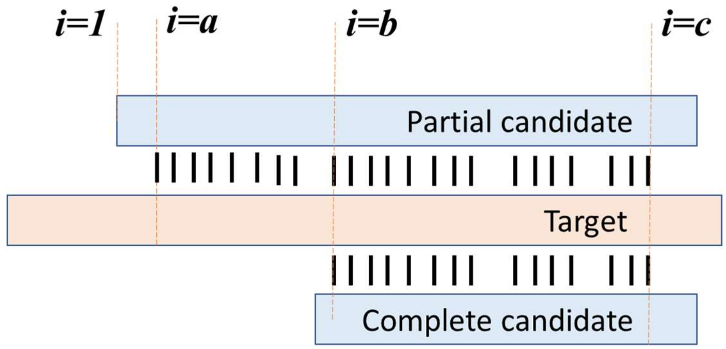


**Supplemental Figure S11**. Illustration for the case when Condition S1 is fulfilled, and the GeneMarkS-T gene prediction is classified as a 5’ *partial CDS*. Here *a, b* and *c* are defined as in Supplemental Fig. S9.

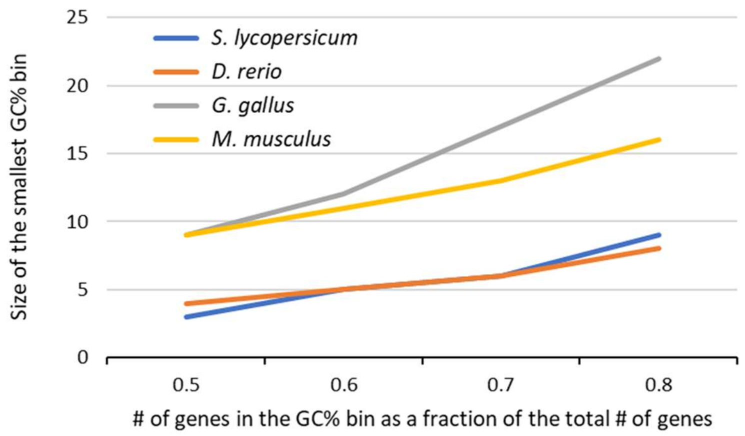


**Supplemental Figure S12**. Results of analysis of the genome GC content inhomogeneity. For each genome, the graphs show the sizes of the narrowest GC% bin in the genome-specific GC content distribution (Y axis) that would contain the number of genes corresponding to a fixed fraction of the total number of annotated genes (X axis). It can be seen from the graph that the *G. gallus* genome is the most GC heterogeneous, followed by the *M. musculus* genome. The remaining two genomes are GC homogeneous: 80% of the whole gene complement can be placed into the GC bin with 10% width.

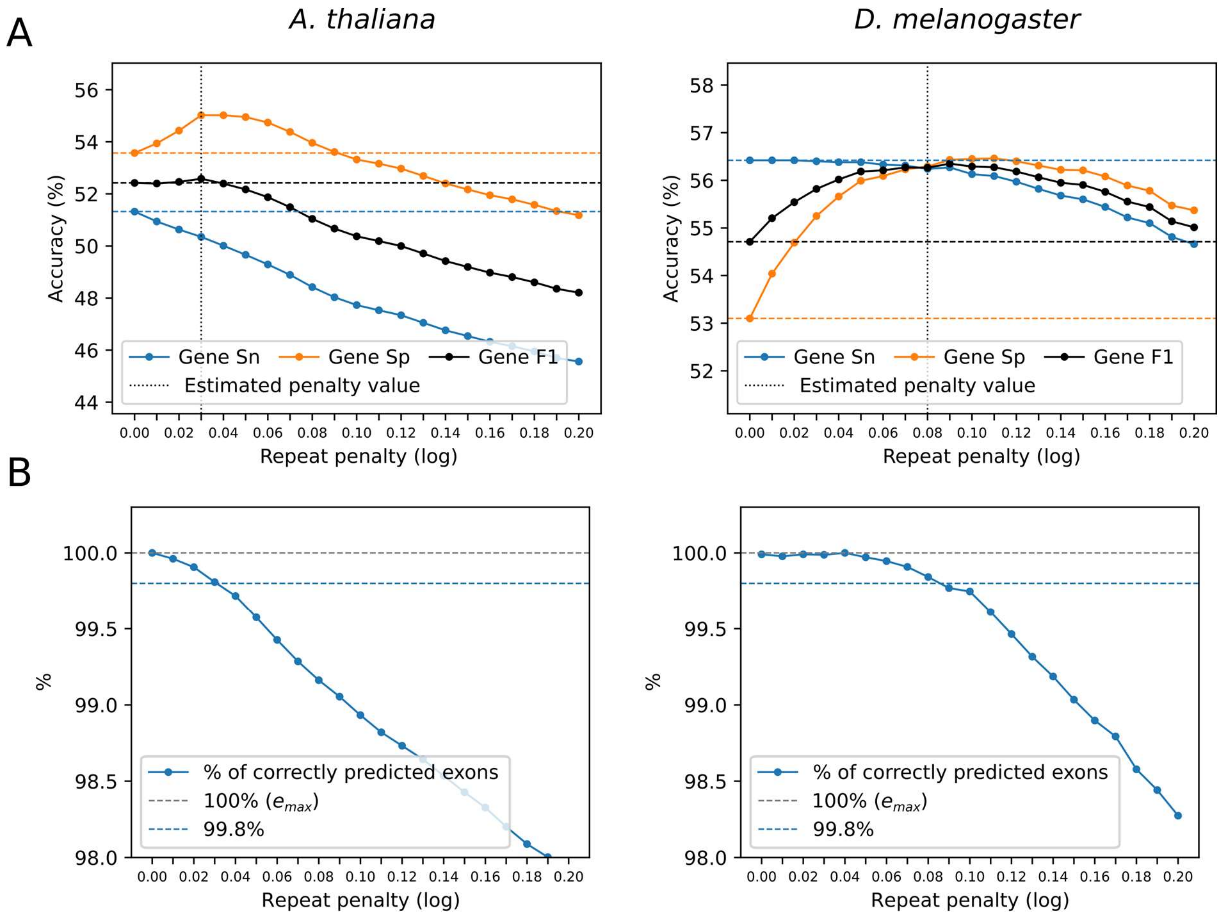


**Supplemental Figure S13. A.** Dependence of the gene level Sn, Pr, and F1 values (determined for the full sets of HC CDSs) on the repeat penalty parameter *q* (natural log) for genomes of *A. thaliana* and *D. melanogaster*. **B.** Dependence of fraction (%) of correctly predicted exons of the HC CDSs (Sn) on the repeat penalty parameter *q* for the same genomes as in A. (See Section S4 of Suppl. Materials)

**Supplemental Table S1**. The numbers of the HC genes predicted by GeneMarkS-TP in the seven genomes, along with Sn/Pr measures computed with respect to the sets of annotated genes as described in the Data Sets section 'Computation of accuracy measures'. The results are listed for the 'Order excluded' and the 'Species excluded' databases (top and bottom sections, respectively).

| Species | Order excluded DB | | Ratio of # of HC genes to # of | Sn/Pr for predicted HC |
|---|---|---|---|---|
| | # of proteins in DB | # of HC genes found | | |
| *C. elegans* | 8,168,321 | 8,062 | 0.40 | 35.7 / 88.4 |
| *A. thaliana* | 3,160,482 | 16,008 | 0.58 | 55.0 / 94.7 |
| *D. melanogaster* | 1,785,203 | 8,109 | 0.58 | 59.6 / 81.8 |
| *S. lycopersicum* | 3,116,328 | 17,231 | 0.68 | 74.9 / 95.2 |
| *D. rerio* | 4,791,893 | 16,918 | 0.66 | 67.0 / 88.5 |
| *G. gallus* | 4,933,362 | 12,473 | 0.72 | 74.4 / 89.1 |
| *M. musculus* | 1,835,426 | 13,057 | 0.58 | 63.5 / 93.2 |
| Species | Species excluded DB | | Ratio of # of HC genes to # of | Sn/Pr for predicted HC |
| | # of proteins in DB | # of HC genes found | | |
| *C. elegans* | 8,245,445 | 11,399 | 0.57 | 51.7 / 90.6 |
| *A. thaliana* | 3,483,291 | 16,551 | 0.60 | 58.8 / 97.6 |
| *D. melanogaster* | 2,588,444 | 9,223 | 0.66 | 63.7 / 96.3 |
| *S. lycopersicum* | 3,456,742 | 17,489 | 0.70 | 75.8 / 95.1 |
| *D. rerio* | 4,973,735 | 16,573 | 0.65 | 66.9 / 90.4 |
| *G. gallus* | 4,984,020 | 12,564 | 0.73 | 74.0 / 88.4 |
| *M. musculus* | 2,228,727 | 12,965 | 0.57 | 63.9 / 94.5 |

**Supplemental Table S2**. Gene and exon level prediction accuracy of the HC CDSs candidates and GeneMark-ETP final gene predictions for the four large genomes (longer than 300 Mbp). The larger F1 values are shown in bold. The gene candidates with no extrinsic match do not make it into the list of genes predicted by GeneMark-ETP.

| Species | | Order excluded DB | | Species excluded DB | |
|---|---|---|---|---|---|
| | | Candidates | Output | Candidates | Output |
| *S. lycopersicum* | gene Sn | 89.5 | 88.2 | 90.6 | 90.2 |
| | gene Pr | 70.6 | 81.4 | 70.9 | 79.8 |
| | gene F1 | 78.9 | **84.7** | 79.5 | **84.6** |
| | exon Sn | 97.1 | 96.7 | 97.4 | 97.2 |
| | exon Pr | 87.1 | 92.6 | 87.0 | 91.6 |
| | exon F1 | 91.8 | **94.6** | 91.9 | **94.3** |
| *D. rerio* | gene Sn | 72.9 | 72.7 | 73.8 | 73.8 |
| | gene Pr | 39.4 | 56.5 | 40.3 | 56.8 |
| | gene F1 | 51.2 | **63.6** | 52.2 | **64.2** |
| | exon Sn | 93.9 | 93.6 | 94.2 | 94.0 |
| | exon Pr | 73.7 | 85.1 | 74.1 | 85.1 |
| | exon F1 | 82.5 | **89.2** | 82.9 | **89.3** |
| *G. gallus* | gene Sn | 78.1 | 78.0 | 77.5 | 77.5 |
| | gene Pr | 40.7 | 67.2 | 40.0 | 65.9 |
| | gene F1 | 53.5 | **72.2** | 52.8 | **71.2** |
| | exon Sn | 95.5 | 95.4 | 95.4 | 95.4 |
| | exon Pr | 76.9 | 90.7 | 76.5 | 90.3 |
| | exon F1 | 85.2 | **93.0** | 85.0 | **92.8** |
| *M. musculus* | gene Sn | 71.7 | 71.3 | 72.8 | 72.7 |
| | gene Pr | 34.5 | 66.0 | 35.3 | 65.9 |
| | gene F1 | 46.5 | **68.6** | 47.6 | **69.1** |
| | exon Sn | 91.6 | 91.2 | 92.0 | 91.7 |
| | exon Pr | 70.1 | 90.7 | 70.7 | 90.7 |
| | exon F1 | 79.4 | **91.0** | 79.9 | **91.2** |

**Supplemental Table S3**. Gene- and exon-level accuracy of the CDSs predictions made by the ab initio GeneMark-ES (ES), the RNA-seq supported GeneMark-ET(ET), the protein similarity supported GeneMark-EP+ (EP+), and GeneMark-ETP (ETP), supported by both extrinsic sources. Genomes longer than 300Mb require extrinsic information to guide the convergence of GeneMark-ES. Therefore, the GeneMark-ES predictions for such genomes were not generated.

| Species | | ES | ET | Order excluded DB | | Species excluded DB | |
|---|---|---|---|---|---|---|---|
| | | | | EP+ | ETP | EP+ | ETP |
| *C. elegans* | Gene Sn | 48.2 | 48.9 | 48.5 | **60.4** | 55.2 | **68.4** |
| | Gene Pr | 47.9 | 48.8 | 46.8 | **67.7** | 53.8 | **73.8** |
| | Gene F1 | 48.0 | 48.8 | 47.6 | **63.8** | 54.5 | **71.0** |
| | Exon Sn | 81.8 | 81.7 | 81.1 | **82.9** | 83.3 | **85.9** |
| | Exon Pr | 83.1 | 83.7 | 82.0 | **90.1** | 84.9 | **91.4** |
| | Exon F1 | 82.5 | 82.7 | 81.5 | **86.4** | 84.1 | **88.6** |
| *A. thaliana* | Gene Sn | 55.8 | 57.1 | 66.6 | **75.8** | 73.4 | **77.9** |
| | Gene Pr | 55.9 | 57.3 | 65.9 | **80.0** | 71.5 | **81.0** |
| | Gene F1 | 55.9 | 57.2 | 66.3 | **77.8** | 72.4 | **79.4** |
| | Exon Sn | 76.9 | 77.1 | 79.8 | **82.3** | 81.5 | **82.9** |
| | Exon Pr | 80.8 | 82.1 | 84.9 | **90.9** | 86.3 | **91.0** |
| | Exon F1 | 78.8 | 79.5 | 82.3 | **86.4** | 83.8 | **86.8** |
| *D. melanogast* | Gene Sn | 51.2 | 53.3 | 56.5 | **71.5** | 69.9 | **78.9** |
| | Gene Pr | 48.5 | 49.7 | 53.9 | **77.9** | 63.5 | **83.1** |
| | Gene F1 | 49.8 | 51.4 | 55.1 | **74.6** | 66.5 | **80.9** |
| | Exon Sn | 67.8 | 68.6 | 70.2 | **76.4** | 76.5 | **80.7** |
| | Exon Pr | 72.8 | 74.2 | 77.3 | **89.7** | 81.1 | **91.4** |
| | Exon F1 | 70.2 | 71.3 | 73.6 | **82.5** | 78.8 | **85.7** |
| *S. lycopersicun* | Gene Sn | | 47.2 | 67.0 | **88.2** | 72.7 | **90.2** |
| | Gene Pr | | 37.4 | 51.3 | **81.4** | 54.8 | **79.8** |
| | Gene F1 | | 41.7 | 58.1 | **84.7** | 62.5 | **84.6** |
| | Exon Sn | | 83.5 | 90.5 | **96.7** | 92.1 | **97.2** |
| | Exon Pr | | 74.2 | 80.0 | **92.6** | 80.7 | **91.6** |
| | Exon F1 | | 78.6 | 84.9 | **94.6** | 86.0 | **94.3** |
| *D. rerio* | Gene Sn | | 20.4 | 35.7 | **72.7** | 39.6 | **73.8** |
| | Gene Pr | | 7.5 | 13.3 | **56.5** | 14.7 | **56.8** |
| | Gene F1 | | 11.0 | 19.4 | **63.6** | 21.4 | **64.2** |
| | Exon Sn | | 79.1 | 84.9 | **93.6** | 86.2 | **94.0** |
| | Exon Pr | | 50.3 | 55.9 | **85.1** | 56.5 | **85.1** |
| | Exon F1 | | 61.5 | 67.4 | **89.2** | 68.2 | **89.3** |
| *G. gallus* | Gene Sn | | 2.4 | 14.1 | **78.0** | 14.4 | **77.5** |
| | Gene Pr | | 1.4 | 11.3 | **67.2** | 11.6 | **65.9** |
| | Gene F1 | | 1.8 | 12.6 | **72.2** | 12.9 | **71.2** |
| | Exon Sn | | 15.1 | 28.7 | **95.4** | 29.0 | **95.4** |
| | Exon Pr | | 27.0 | 53.4 | **90.7** | 53.8 | **90.3** |
| | Exon F1 | | 19.3 | 37.3 | **93.0** | 37.7 | **92.8** |
| *M. musculus* | Gene Sn | | 7.8 | 22.0 | **71.3** | 23.7 | **72.7** |
| | Gene Pr | | 5.4 | 15.0 | **66.0** | 16.0 | **65.9** |
| | Gene F1 | | 6.4 | 17.8 | **68.6** | 19.1 | **69.1** |
| | Exon Sn | | 49.7 | 57.3 | **91.2** | 58.1 | **91.7** |
| | Exon Pr | | 50.9 | 64.2 | **90.7** | 64.8 | **90.7** |
| | Exon F1 | | 50.3 | 60.6 | **91.0** | 61.3 | **91.2** |

**Supplemental Table S4**. Comparison of gene- and exon-level accuracy of CDS predictions made by RNA-seq-based BRAKER1, protein-based BRAKER2, TSEBRA, and GeneMark-ETP (ETP). Note that the low accuracy in genomes of *G. gallus* and *M. musculus* observed for BRAKER1, BRAKER2, and TSEBRA could be related to the use of a single statistical model for the genome-wide CDS prediction. GeneMark-ETP uses several GC-specific models.

| Species | | BRAKER1 | Order excluded DB | | | Species excluded DB | | |
|---|---|---|---|---|---|---|---|---|
| | | | BRAKER2 | TSEBRA | ETP | BRAKER2 | TSEBRA | ETP |
| *C. elegans* | Gene Sn | **61.8** | 46.8 | 60.3 | 60.4 | 69.0 | **71.1** | 68.4 |
| | Gene Pr | 65.6 | 54.1 | **77.5** | 67.7 | 70.1 | **80.5** | 73.8 |
| | Gene F1 | 63.6 | 50.2 | **67.8** | 63.8 | 69.6 | **75.5** | 71.0 |
| | Exon Sn | **85.0** | 74.0 | 76.6 | 82.9 | 84.8 | 83.9 | **85.9** |
| | Exon Pr | 88.5 | 87.8 | **93.4** | 90.1 | 91.5 | **93.8** | 91.4 |
| | Exon F1 | **86.7** | 80.3 | 84.2 | 86.4 | 88.0 | **88.6** | **88.6** |
| *A. thaliana* | Gene Sn | 59.6 | 72.6 | 73.6 | **75.8** | 79.2 | **79.3** | 77.9 |
| | Gene Pr | 61.3 | 70.1 | **81.2** | 80.0 | 75.6 | **82.8** | 81.0 |
| | Gene F1 | 60.4 | 71.3 | 77.2 | **77.8** | 77.4 | **81.0** | 79.4 |
| | Exon Sn | 78.3 | 81.0 | 79.6 | **82.3** | 83.1 | 82.7 | **82.9** |
| | Exon Pr | 82.5 | 88.4 | **93.7** | 90.9 | 88.2 | **93.2** | 91.0 |
| | Exon F1 | 80.4 | 84.5 | 86.1 | **86.4** | 85.6 | **87.6** | 86.8 |
| *D. melanogaster* | Gene Sn | 63.8 | 61.1 | 68.0 | **71.5** | 78.9 | **80.0** | 78.9 |
| | Gene Pr | 62.3 | 60.9 | 75.4 | **77.9** | 73.6 | 80.9 | **83.1** |
| | Gene F1 | 63.0 | 61.0 | 71.5 | **74.6** | 76.1 | 80.4 | **80.9** |
| | Exon Sn | **77.0** | 71.4 | 72.1 | 76.4 | 80.1 | 79.8 | **80.7** |
| | Exon Pr | 80.9 | 83.4 | **89.9** | 89.7 | 88.5 | **92.2** | 91.4 |
| | Exon F1 | 78.9 | 76.9 | 80.0 | **82.5** | 84.1 | 85.6 | **85.7** |
| *S. lycopersicum* | Gene Sn | 61.8 | 79.6 | 82.5 | **88.2** | 84.2 | 85.4 | **90.2** |
| | Gene Pr | 47.1 | 56.5 | 71.3 | **81.4** | 58.9 | 72.1 | **79.8** |
| | Gene F1 | 53.5 | 66.1 | 76.5 | **84.7** | 69.3 | 78.2 | **84.6** |
| | Exon Sn | 90.7 | 94.2 | 94.9 | **96.7** | 95.4 | 96.1 | **97.2** |
| | Exon Pr | 75.5 | 82.8 | 90.3 | **92.6** | 82.3 | 90.2 | **91.6** |
| | Exon F1 | 82.4 | 88.1 | 92.5 | **94.6** | 88.4 | 93.0 | **94.3** |
| *D. rerio* | Gene Sn | 51.7 | 55.0 | 66.9 | **72.7** | 57.8 | 69.0 | **73.8** |
| | Gene Pr | 28.1 | 29.5 | 45.7 | **56.5** | 27.9 | 46.0 | **56.8** |
| | Gene F1 | 36.4 | 38.4 | 54.3 | **63.6** | 37.6 | 55.2 | **64.2** |
| | Exon Sn | 91.1 | 88.0 | 89.4 | **93.6** | 89.4 | 90.1 | **94.0** |
| | Exon Pr | 75.4 | 78.9 | **87.2** | 85.1 | 76.2 | **86.8** | 85.1 |
| | Exon F1 | 82.5 | 83.2 | 88.3 | **89.2** | 82.2 | 88.4 | **89.3** |
| *G. gallus* | Gene Sn | 6.6 | 25.2 | 26.7 | **78.0** | 27.2 | 28.3 | **77.5** |
| | Gene Pr | 3.5 | 16.6 | 22.2 | **67.2** | 18.1 | 23.3 | **65.9** |
| | Gene F1 | 4.6 | 20.0 | 24.2 | **72.2** | 21.7 | 25.6 | **71.2** |
| | Exon Sn | 66.1 | 35.0 | 59.8 | **95.4** | 35.3 | 60.0 | **95.4** |
| | Exon Pr | 48.1 | 59.2 | 74.4 | **90.7** | 60.6 | 74.4 | **90.3** |
| | Exon F1 | 55.7 | 44.0 | 66.3 | **93.0** | 44.6 | 66.4 | **92.8** |
| *M. musculus* | Gene Sn | 27.8 | 32.5 | 44.2 | **71.3** | 35.9 | 46.7 | **72.7** |
| | Gene Pr | 14.8 | 21.2 | 31.3 | **66.0** | 23.2 | 32.7 | **65.9** |
| | Gene F1 | 19.3 | 25.7 | 36.7 | **68.6** | 28.2 | 38.5 | **69.1** |
| | Exon Sn | 83.9 | 57.6 | 77.4 | **91.2** | 59.3 | 78.1 | **91.7** |
| | Exon Pr | 67.5 | 71.6 | 83.3 | **90.7** | 72.7 | 83.5 | **90.7** |
| | Exon F1 | 74.8 | 63.8 | 80.2 | **91.0** | 65.3 | 80.7 | **91.2** |

**Supplemental Table S5**. The CDS prediction accuracy of MAKER2 and GeneMark-ETP was assessed for the three model species. The results given for MAKER2 are supposed to be upper bounds reached among possible methods of MAKER2 training. The protein databases are described in Section S3 of Supplemental Materials.

| | | *D. melanogaster* | | *D. rerio* | | *M. musculus* | |
|---|---|---|---|---|---|---|---|
| | | MAKER2 | ETP | MAKER2 | ETP | MAKER2 | ETP |
| exon | Sn | 75.2 | **80.7** | 83.3 | **93.9** | 79.2 | **91.7** |
| | Pr | 74.0 | **91.4** | 79.2 | **84.9** | 77.4 | **87.9** |
| | F1 | 74.6 | **85.7** | 81.2 | **89.2** | 78.3 | **89.8** |
| gene | Sn | 60.2 | **79.0** | 47.7 | **73.5** | 41.6 | **73.1** |
| | Pr | 55.3 | **83.0** | 37.6 | **56.2** | 34.8 | **59.7** |
| | F1 | 57.7 | **81.0** | 42.0 | **63.7** | 37.9 | **65.7** |

**Supplemental Table S6**. The BUSCO scores for the GeneMark-ETP predictions made with the 'Order excluded' databases and for genes annotated in RefSeq. If a protein-coding gene had alternative CDSs, the longest CDS was selected for the analysis. The BUSCO database version odb10 and the BUSCO software version 5.5.0 were used.

| Species | BUSCO database lineage name | Number of gene models in the BUSCO database | BUSCO score for GeneMark-ETP gene predictions | BUSCO score for the genes annotated in RefSeq |
|---|---|---|---|---|
| *C. elegans* | *nematoda* | 3,131 | 98.9% | 100.0% |
| *A. thaliana* | *brassicales* | 4,596 | 99.0% | 99.9% |
| *D. melanogaster* | *diptera* | 3,285 | 97.2% | 99.8% |
| *S. lycopersicum* | *solanales* | 5,950 | 98.2% | 99.0% |
| *D. rerio* | *actinopterygii* | 3,640 | 94.8% | 98.7% |
| *G. gallus* | *aves* | 8,338 | 95.9% | 98.0% |
| *M. musculus* | *glires* | 13,798 | 92.6% | 99.5% |

**Supplemental Table S7**. Genomic sequences (assembly versions) and genome annotations that were used in the computational experiments. Additional details for *C. elegance*, *D. melanogaster*, and *S. lycopersicum* annotations are as follows. Both NCBI and Ensemble adapted *C. elegance* annotation from Wormbase. We specifically used the Wormbase version WS284 (Feb 2022). Both NCBI and Ensemble adapted *D. melanogaster* annotation from Flybase. We specifically used the Flybase version r6.44 (Feb 2022). Ensemble adapts the annotation from the International Tomato Annotation Group. We used the annotation version ITAG3.2 (Jun 2017). A date in parenthesis is the date of the last update prior to the data use.

| Species | Assembly version | Annotation 1 | Annotation 2 |
|---|---|---|---|
| *C. elegans* | GCF_000002985.6 | NCBI RefSeq (adopted from Wormbase) | Ensembl (adopted from Wormbase) |
| *A. thaliana* | GCF_000001735.4 | NCBI RefSeq (adopted from Araport11) | EnsemblPlants (adopted from Araport11) |
| *D. melanogaster* | GCF_000001215.4 | NCBI RefSeq (adopted from FlyBase) | Ensembl (adopted from FlyBase) |
| *S. lycopersicum* | GCF_000188115.4 | NCBI RefSeq release 103 | Ensemble (adopted from ITAG) |
| *D. rerio* | GCF_000002035.6 | NCBI RefSeq release 106 | Ensembl GRCz11.105 |
| *G. gallus* | GCF_000002315.6 | NCBI RefSeq release 104 | Ensembl GRCg6a.105 |
| *M. musculus* | GCF_000001635.27 | NCBI RefSeq release 109 | Ensembl release 109 |

**Supplemental Table S8**. Selection of species from OrthoDB v10.1. Numbers in bold black font show the number of species in the largest OrthoDB segment, IP0, considered for a given species. Numbers in bold blue font show the number of species excluded from IP0 in the 'Order excluded' segment of OrthoDB. The 'Species excluded' segment of OrthoDB comprises all proteins in IP0 but those from the species of interest itself.

| Species | # of species in the OrthoDB clade | | | | | | Name of the OrthoDB segment | # of proteins in the OrthoDB segment (M) |
|---|---|---|---|---|---|---|---|---|
| | Genus | Family | Order | Class | Phylum | Kingdom | | |
| *C. elegans* | 3 | 3 | **5** | 6 | 7 | **448** | *Metazoa* | 8.3 |
| *A. thaliana* | 2 | 8 | **10** | - | 100 | **117** | *Plantae* | 3.5 |
| *D. melanogaster* | 20 | 20 | **56** | 148 | **170** | - | *Arthropoda* | 2.6 |
| *S. lycopersicum* | 2 | 10 | **11** | - | 100 | **117** | *Plantae* | 3.5 |
| *D. rerio* | 1 | 5 | **5** | 50 | **246** | - | *Chordata* | 5.0 |
| *G. gallus* | 1 | 3 | **4** | 62 | **246** | - | *Chordata* | 5.0 |
| *M. musculus* | 3 | 5 | **20** | **111** | - | - | *Mammalia* | 2.3 |

**Supplemental Table S9**. The RNA-seq libraries used for the computational experiments.

| Species | RNA-Seq library ID | Number of paired reads (M) | Read length (nt) | Library size (Gb) |
|---|---|---|---|---|
| *C. elegans* | SRR065717 | 29.1 | 76 | 4.4 |
| | SRR065719 | 73.3 | 76 | 11.1 |
| | SRR473298 | 19.9 | 100 | 4.0 |
| | SRR2054452 | 10.2 | 100 | 2.0 |
| | Total | 132.5 | | 21.5 |
| *A.thaliana* | SRR934391 | 20.0 | 101 | 4.0 |
| | SRR5588566 | 24.7 | 125 | 6.2 |
| | SRR7169927 | 19.2 | 101 | 3.9 |
| | Total | 63.9 | | 14.1 |
| *D. melanogaster* | SRR023505 | 8.4 | 76 | 1.3 |
| | SRR023546 | 8.9 | 76 | 1.4 |
| | SRR023608 | 11.9 | 76 | 1.8 |
| | SRR026433 | 22.1 | 76 | 3.4 |
| | SRR027108 | 7.2 | 76 | 1.1 |
| | Total | 58.5 | | 9.0 |
| *S. lycopersicum* | SRR2002284 | 56.2 | 73 | 8.2 |
| | SRR7959012 | 25.4 | 149 | 7.6 |
| | SRR7959019 | 27.9 | 149 | 8.3 |
| | SRR14055940 | 21.2 | 150 | 6.4 |
| | Total | 130.7 | | 30.5 |
| *D. rerio* | SRR9735169 | 28.2 | 75 | 4.2 |
| | SRR10004226 | 21.6 | 150 | 6.5 |
| | SRR10040127 | 25.9 | 126 | 6.5 |
| | Total | 75.7 | | 17.2 |
| *G. gallus* | ERR2812450 | 44.9 | 150 | 13.5 |
| | SRR3971633 | 24.0 | 100 | 4.8 |
| | SRR6337028 | 10.0 | 100 | 2.0 |
| | SRR11038071 | 16.4 | 151 | 5.0 |
| | Total | 95.3 | | 25.3 |
| *M. musculus* | SRR567480 | 155.7 | 101 | 31.5 |
| | SRR567482 | 161.1 | 101 | 32.5 |
| | SRR567497 | 94.3 | 101 | 19.0 |
| | Total | 411.1 | | 83.0 |

**Supplemental Table S10**. The Sn and Pr of the predicted HC CDSs, complete and partial, are shown for the 'Order excluded' and for the 'Species excluded' databases (see Data Sets). The values of Sn and Pr are determined for i/ a set of initial CDSs predictions inferred from GeneMarkS-T predictions made in transcripts and ii/ a set of high-confidence (HC) CDSs, the output of GeneMarkS-TP. The Sn and Pr are shown separately for the complete and partial GeneMarkS-T predictions and for complete and partial HC CDSs. The true positive prediction of a partial CDS is called if the partial prediction coincides with a part of a CDS in the reference annotation. A significant increase in the Pr values of the partial CDSs occurred in the transition from the initial GeneMarkS-T predictions to the HC CDSs.

| Species | | GeneMarkS-T predictions | | HC genes (processed by GeneMarkS-TP) | | | |
|---|---|---|---|---|---|---|---|
| | | | | Order excluded DB | | Species excluded DB | |
| | | Complete | Partial | Complete | Partial | Complete | Partial |
| *C. elegans* | Sn | 42.9 | 3.9 | 33.6 | 2.1 | 47.7 | 4.0 |
| | Pr | 82.0 | 18.2 | 88.8 | 81.5 | 91.5 | 80.7 |
| *A. thaliana* | Sn | 49.8 | 1.4 | 55.6 | 1.1 | 57.3 | 1.6 |
| | Pr | 89.1 | 17.0 | 97.4 | 92.3 | 97.8 | 90.8 |
| *D. melanogaster* | Sn | 56.4 | 3.2 | 53.3 | 1.8 | 60.6 | 3.1 |
| | Pr | 87.5 | 38.1 | 95.0 | 85.3 | 96.9 | 85.0 |
| *S. lycopersicum* | Sn | 66.3 | 1.4 | 73.7 | 1.3 | 74.2 | 1.5 |
| | Pr | 84.1 | 26.6 | 95.4 | 87.2 | 95.4 | 84.8 |
| *D. rerio* | Sn | 55.3 | 4.3 | 62.8 | 4.2 | 62.4 | 4.5 |
| | Pr | 68.4 | 32.8 | 89.7 | 78.9 | 92.8 | 75.3 |
| *G. gallus* | Sn | 43.9 | 5.7 | 67.9 | 6.5 | 66.3 | 7.7 |
| | Pr | 64.0 | 23.0 | 89.5 | 86.1 | 90.0 | 80.3 |
| *M. musculus* | Sn | 48.4 | 1.2 | 60.8 | 2.7 | 60.5 | 3.4 |
| | Pr | 80.4 | 9.6 | 95.1 | 68.0 | 96.7 | 69.8 |

**Supplemental Table S11**. The values of the genome-specific masking penalty parameter q (in natural logarithms). The optimal q value was determined for each GC bin for GC-heterogeneous genomes.

| | Order excluded DB | | | Species excluded DB | | |
|---|---|---|---|---|---|---|
| *C. elegans* | 0.06 | | | 0.05 | | |
| *A. thaliana* | 0.03 | | | 0.03 | | |
| *D. melanogaster* | 0.08 | | | 0.08 | | |
| *S. lycopersicum* | 0.04 | | | 0.04 | | |
| *D. rerio* | 0.08 | | | 0.09 | | |
| *GC* | Low | Medium | High | Low | Medium | High |
| *G. gallus* | 0.15 | 0.17 | 0.12 | 0.14 | 0.16 | 0.11 |
| *M. musculus* | 0.13 | 0.14 | 0.14 | 0.14 | 0.14 | 0.14 |

### References


Buchfink B, Xie C, Huson DH. 2015. Fast and sensitive protein alignment using DIAMOND. *Nature Methods* **12**: 59-60.

Kirkpatrick S, Gelatt CD, Vecchi MP. 1983. Optimization by Simulated Annealing. *Science* **220**: 671-680.

Kozak M. 1987. An analysis of 5'-noncoding sequences from 699 vertebrate messenger RNAs. *Nucleic Acids Res* **15**: 8125-8148.

Kozak M. 1999. Initiation of translation in prokaryotes and eukaryotes. *Gene* **234**: 187-208.

Tang S, Lomsadze A, Borodovsky M. 2015. Identification of protein coding regions in RNA transcripts. *Nucleic Acids Res* **43**: e78.

Zhao W, He X, Hoadley KA, Parker JS, Hayes DN, Perou CM. 2014. Comparison of RNA-Seq by poly (A) capture, ribosomal RNA depletion, and DNA microarray for expression profiling. *BMC Genomics* **15**: 419.
